# Cell-free quorum sensing to drive feedback and communication within populations of droplet interface bilayers

**DOI:** 10.1101/2023.03.04.531082

**Authors:** David T. Gonzales, Surased Suraritdechachai, Christoph Zechner, T-Y Dora Tang

**Affiliations:** Max Planck Institute of Molecular Cell Biology and Genetics, 01307 Dresden, Germany; Center for Systems Biology Dresden, 01307 Dresden, Germany; Physics of Life, Cluster of Excellence, TU Dresden, 01603 Dresden, Germany

## Abstract

Building synthetic multicellular systems using non-living molecular components is a grand challenge in the field of bottom-up synthetic biology. Towards this goal, a diverse range of chemistries have been developed to provide mechanisms of intercellular communication and methods to assemble multicellular compartments. However, building bottom-up synthetic multicellular systems is still challenging because it requires the integration of intercellular reaction networks with compatible cellular compartment properties. In this study, we encapsulated cell-free expression systems (CFES) expressing two quorum sensing genetic circuits into droplet interface bilayer (DIB) synthetic cells to demonstrate intercellular communication and feedback. We further develop a method of generating custom DIB multicellular structures by acoustic liquid handling to automatically dispense the CFES droplets and show the potential for multiplexing compartmentalized gene circuits for generating heterogeneous populations of cells. Our work provides a step towards building more complex multicellular systems with feedback mechanisms from the bottom-up to study and experimentally model biological multiscalar processes.

## 1. Introduction

One of the grand challenges in synthetic biology is to build and control synthetic cell populations with increasing levels of molecular complexity. In biology, multicellular systems use communication to co-ordinate cells across length scales and to drive high levels of functionality such as in quorum sensing microbial communities [1], wound healing in tissues [2], and cell fate decisions during differentiation [3]. Thus, integrating communication between synthetic cells is essential to build multicellular systems [4]. Bottom-up synthetic biology offers a diverse range of tools to incorporate communication within synthetic cells. Encapsulated chemical reaction networks can provide the intracellular machinery of the synthetic cell, whilst the boundary conditions provided by micron-sized compartments allow the selective diffusion of molecules to act as intercellular signaling molecules within the reaction network. This concept has been demonstrated using lipid vesicles with membrane pore proteins encapsulating cell-free expression systems (CFES) [5] or enzyme cascades [6,7], droplet interface bilayers (DIB) with CFES [8], porous clay-DNA hydrogel cell mimics with CFES [9], and proteinosomes with DNA strand displacement (DSD) [10,11] or PEN DNA reactions [12]. In addition, depending on the diffusion properties of the signaling molecule and the structure of the cell population, communication can occur directly between cells in contact [8] or through the surrounding medium into which the signal molecule is released [5,6]. Importantly, intercellular communication combined with the modular nature of compartmentalized reactions allows us to further implement complex and multiscale behavior. For instance, feedback loops between activated particles [13] or populations of proteinosomes encapsulating DSD [10] or PEN DNA reactions [12], and DIBs with CFES [14] have been shown to result in travelling waves, stochastic differentiation, population-wide transient activation and autocatalysis. Not only has the implementation of intercellular communication demonstrated our increasing control over synthetic systems, communication also provides a viable route to integrate synthetic cellular systems with biological cells [15]. In addition, communicating synthetic cellular populations offer model systems to understand the role of spatial organization, compartmentalization, and signal diffusion properties in multicellular systems [11,16].

In the present study, we incorporated CFES and DIBs to generate synthetic cells capable of cell-to-cell communication and feedback using quorum sensing genetic circuits. Quorum sensing (QS) communication, first described in the bioluminescent marine bacterium *Vibrio fischeri*, is a form of secrete-and-sensing communication where each cell secretes a membrane-diffusible signaling molecule that diffuses to the external environment where it can be detected by other cells within the population [1]. Gene circuits based on QS systems are one of the most widely used forms of cell-to-cell communication in synthetic biology and have been well-established to function in *E. coli* extract-based CFE reactions [17,18]. QS genetic circuits in CFE reactions have also been encapsulated into liposomes [15] and in DIBs [14,19]. However, to date the implementation of feedback mechanisms based on the QS systems have not yet been demonstrated. Achieving this offers a novel way to co-ordinate population level behavior via the regulation of gene circuits in cell free conditions. To this end, we aimed to generate synthetic cellular populations capable of cell-to-cell communication and feedback using QS genetic circuits under cell free environments. We used two QS systems from *V. fischeri* (Lux QS) and *Pseudomonas aeruginosa* (Las QS), which are each composed of three genetic parts: *luxI, luxR*, and *pLux* for the Lux QS system and *lasI, lasR*, and *pLas* for the Las QS system (Fig. 1A and B). The *luxI* and *lasI* genes encode for the enzymes - acyl-homoserine lactone (acyl-HSL) synthases that produce the signalling molecule, N-(3-oxohexanoyl)homoserine lactone (3OC6-HSL) and N-(3-oxododecanoyl)homoserine lactone (3OC12-HSL) for their respective QS systems (Fig. 1A and B). By utilizing an *E. coli* extract-based CFES, we circumvented the need to externally supply the precursors (acyl-acyl carrier protein and S-adenosyl methionine) required for acyl-HSL signaling molecule synthesis [17]. The *luxR* and *lasR* genes encode the transcription factors that bind with their respective acyl-HSL signaling molecules and induce expression by the promoters of the *lux* and *las* operon (*pLux* and *pLas*). These gene circuits were encapsulated into DIBs to act as communicating synthetic cells. DIBs are water-in-oil droplets stabilized by phospholipids which spontaneously form a lipid monolayer at the interface between the water and oil phases [20]. When two aqueous droplets are brought in contact together, a lipid bilayer is formed at the contact surface between the two droplets. This maintains the separation of the contents of the two droplets but allows the acyl-HSL signal molecules to diffuse across the bilayer. We show that the gene circuits within the DIBs sense and respond to each other to form intercellular communication and feedback loops. Lastly, inspired by recent advances in 3D-droplet printing [8,21], we then developed a convenient method to generate custom and uniform-sized populations of DIB synthetic cells using acoustic liquid handling technology. Taken together, our work demonstrates the utility of compartmentalized CFES to support higher-order functions and lays the foundation to develop strategies to co-ordinate dynamic behavior in populations of cells and regulate population heterogeneity.

**Figure 1.**
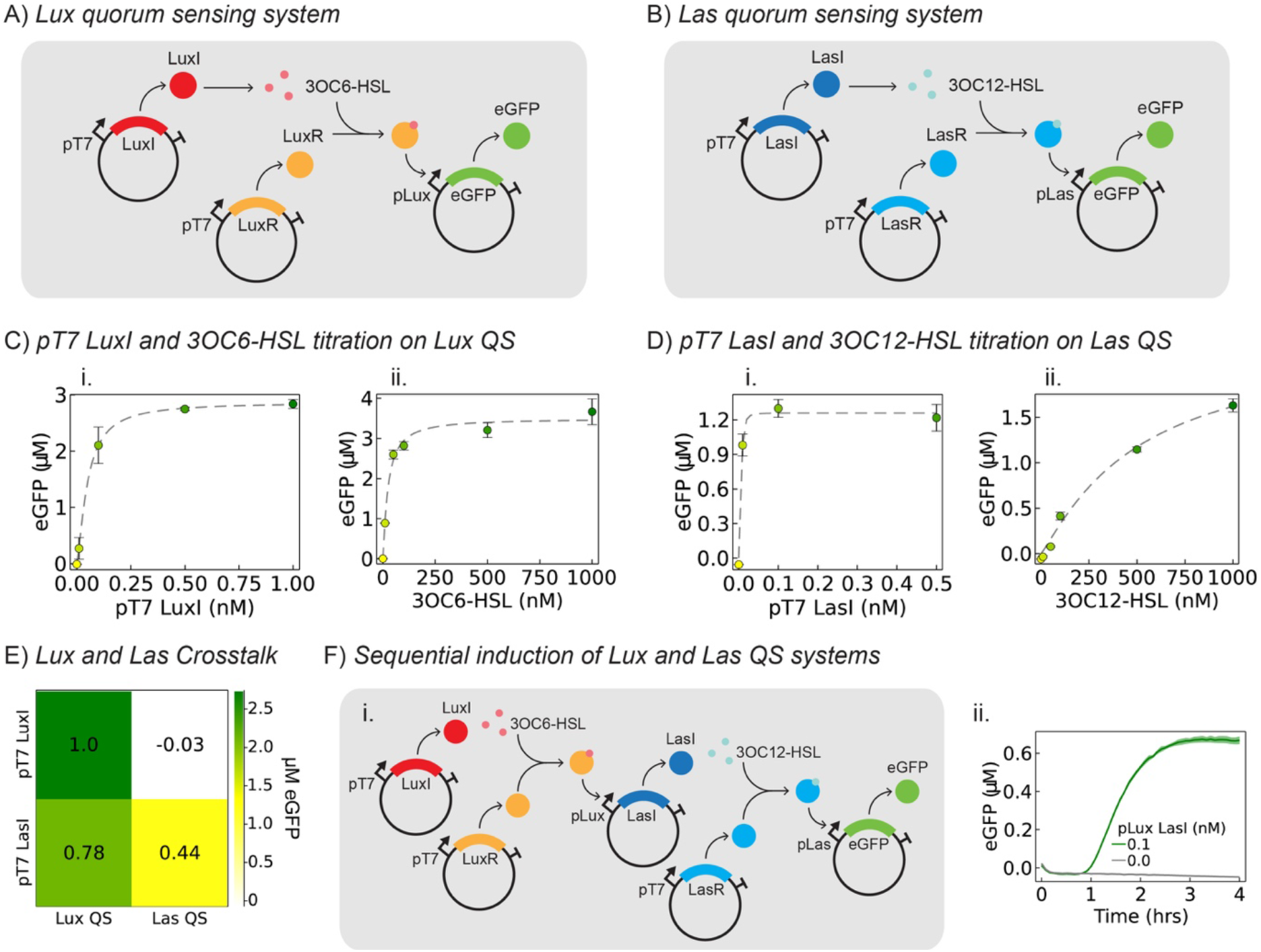
Characterization of Lux and Las quorum sensing gene circuits in bulk CFES. Schematic of **(A)** Lux QS gene circuit plasmids consisting of the sender plasmid (pT7 LuxI) and the receiver plasmids (pT7 LuxR and pLux eGFP) and **(B)** Las QS gene circuit plasmids consisting of the sender plasmid (pT7 LasI) and the receiver plasmids (pT7 LasR and pLas eGFP). **(C)** eGFP expression in Lux QS gene circuit with increasing **(i)** pT7 LuxI plasmid (0-1 nM) and **(ii)** 3OC6-HSL (0-1000 nM) concentration with 0.5 nM pT7 LuxR and 10 nM pLux eGFP. Plots show expressed eGFP concentration after 4 hrs. **(D)** eGFP expression in Las QS gene circuit with increasing **(i)** pT7 LuxI plasmid (0-0.5 nM) and **(ii)** 3OC6-HSL (0-1000 nM) concentration with 0.5 nM pT7 LasR and 10 nM pLas eGFP. Plots show expressed eGFP concentration after 4 hrs. Dashed lines in C and D are Hill function fits. Error bars are the standard deviation from three repeats. **(E)** Characterization of crosstalk between the Lux and Las QS circuits. 0.5 nM pT7 LuxI or 0.5 nM pT7 LasI plasmids. Color of each square indicates the concentration of eGFP expressed at endpoint (t = 4 hours) as noted in the color bar on the right. Values written within each square is the normalized eGFP concentration with respect to the highest value given by the expression of pT7 LuxI. **(F) (i)** Schematic of linear Lux and Las QS circuit. The CFE reaction consists of 0.5 nM pT7 LuxI, 0.5 nM pT7 LuxR, 0.1 nM pLux LasI, 0.5 nM pT7 LasR, and 10 nM pLas eGFP plasmids. **(ii)** Timeseries eGFP expression with and without the intermediate plasmid pLux LasI. All data was obtained using a TECAN Genios Pro plate reader with an incubation temperature of 30°C.

## 2. Results and discussion

### Characterization of quorum sensing gene circuits in bulk CFE reactions

To test the Lux and Las QS gene circuits in cell free conditions, we constructed three separate plasmids for each QS component. A constitutive T7 RNA polymerase (RNAP)-mediated promoter is used to drive the expression of the acyl-HSL synthases (pT7 LuxI and pT7 LasI plasmids) and transcription factors (pT7 LuxR and pT7 LasR plasmids). The signaling molecules 3OC6-HSL/3OC12-HSL produced from expressed LuxI/LasI binds to the transcription factors LuxR/LasR to induce eGFP expression from their corresponding QS promoters in the pLux eGFP and pLas eGFP plasmids.

We first determined the optimized concentrations for each plasmid component using a series of pilot experiments with the Lux and Las QS system plasmids. This was important as the expression level of each component must be balanced with the limited gene expression resources under cell-free conditions. For instance, increasing the pLas eGFP plasmid concentration from 5 nM to 10 nM increased the induced eGFP signal. In contrast, increasing the concentration of pT7 LuxI plasmid above 1nM can reduce the overall induced expression of eGFP as more gene expression resources are used to produce LuxI instead of eGFP (*see Fig. S2 in the Supplementary Materials*). To achieve a high signal from the induced eGFP expression, we set concentrations of 0.5 nM pT7 LuxR and 10 nM pLux eGFP for the Lux QS system and 0.5 nM pT7 LasR and 10 nM pLas eGFP for the Las QS system. We then determined the eGFP expression profiles of the Lux and Las QS systems against increasing concentrations of pT7 LuxI (Fig. 1Ci) or pT7 LasI plasmid (Fig. 1Di). We found that the Lux QS system had a maximal induced expression of approximately 3 μM eGFP at 0.5 nM pT7 LuxI plasmid and the Las QS system at 1.3 μM eGFP at 0.1 nM pT7 LasI plasmid. To confirm that the induction of eGFP was driven by the acyl-HSL signaling molecules, we titrated the acyl-HSL molecules directly into the CFES containing either 0.5 nM pT7 LuxR and 10 nM pLux eGFP (Fig. 1Cii) or 0.5 nM pT7 LasR and 10 nM pLux eGFP (Fig. 1Dii) in the absence of the plasmids expressing acyl-HSL synthases (pT7 LuxI or pT7 LasI). The results show increasing amounts of eGFP expression with increasing concentration of acyl-HSL. This confirms that eGFP expression is induced by acyl-HSL signaling molecules that are produced *in situ* after the expression of acyl-HSL synthases and is consistent with previous work [15,17,18].

Our results also showed, that the dynamic range of the signaling molecule required to induce the QS system is narrower for the Lux QS (3OC6-HSL: 0-250 nM) compared to the Las QS (3OC12-HSL: 0-1000 nM). This was also observed in previous studies in *E. coli* extract-based CFES [22]. In contrast, induction by acyl-HSL synthase expression from the input plasmid concentration of the Las QS system has a narrower range (0-0.1 nM of pT7 LasI) compared to the Lux QS system (0-0.5 nM pT7 LuxI). These differences in reporter protein expression between direct acyl-HSL and acyl-HSL synthase plasmid induction could be attributed to differences in gene expression rate and/or enzymatic conversion rates of the acyl-HSL synthases, which could affect overall yield of eGFP expression in a resource-limited environment. Taken together, our results show that the Lux and Las QS system are active in our *E. coli* extract-based CFES and driven by production and induction of the acyl-HSL signaling molecules. The Lux system has a broader dynamic range, produces more eGFP and requires lower concentrations of signaling molecules for induction compared to the Las system suggesting that the Lux system is more sensitive and efficient compared to the Las system in bulk cell free expression system.

To determine whether the Lux and Las QS systems could be combined together in a gene circuit, we evaluated the degree of cross-talk between the Lux and Las QS systems (Fig. 1E and *Fig. S6 in the Supplementary Materials*). To this end, we added 0.5 nM pT7 LuxI plasmid into the Las QS system (0.5 nM pT7 LasR and 10 nM pLas eGFP) and 0.5 nM pT7 LasI plasmid into the Lux QS system (0.5 nM pT7 LuxR and 10 nM pLux eGFP) in our CFES. Expression of eGFP showed that the Lux QS system was induced by the LasI plasmid to 78% of the expression level by induction with the LuxI plasmid. In comparison, the Las QS system is not induced by the pT7 LuxI plasmid at the plasmid concentrations used. The same pattern of cross-reaction between the Lux and Las QS system in *E. coli* extract-based CFE reactions was also observed by Halleran and Murray (2018) [22]. To determine if this cross-reaction is due to the promiscuity of the acyl-AHL synthases or the binding and induction of the transcription factor:acyl-HSL complex, we performed the same cross-reaction experiments using 3OC6-HSL and 3OC12-HSL to directly induce the QS systems (*Fig. S6 in the Supplementary Materials*). We found that the relative amounts of cross-reaction are comparable when using acyl-HSL synthase plasmids and the acyl-HSLs, which suggests that the cross-reaction of pT7 LasI to the Lux QS system occurs downstream at the level of 3OC12-HSL.

Given that 3OC12-HSL induces both the Lux and Las QS systems and 3OC6-HSL induces only the Lux QS system, we tested a five-plasmid gene circuit where the Lux QS system induces the Las QS system in a sequential order (Fig. 1F). Here, LuxI and LuxR are produced constitutively, LuxI produces 3OC6-HSL that binds to LuxR to induce expression of LasI through the promoter pLux. LasI synthesizes 3OC12-HSL that binds to LasR (expressed constitutively) to induce eGFP expression on the Las promoter. Given that increasing concentrations of pLux LasI can induce leaky expression of the Las systems we used low concentration of pLux LasI (0.1 nM) in this linear gene circuit (*Fig. S7 in the Supplementary Materials*). Positive and negative expression of eGFP in the presence and absence of the intermediate step provided by the pLux LasI plasmid confirms the stepwise induction of the sequential five-plasmid gene circuit in CFES.

### Quorum sensing gene circuits in droplet interface bilayers

Having established and shown successful implementation of a five-plasmid gene circuit in bulk CFE reactions, we next encapsulated the QS gene circuits within droplet interface bilayers. Compartmentalization of the gene circuits provides spatial localization of distinct parts of the circuit which can be connected by molecular diffusion of a signaling molecule. Not only could this provide communication strategies across populations, it can also change the functionality of the gene circuit. In this case the functionality can be switched from a sequential induction in bulk solution to a closed-loop feedback mechanism within two communicating cells.

To this end, the Lux and Las QS plasmids were incorporated into DIBs by manually pipetting ∼0.05-0.1 μL of the CFES for cell 1 and 2 directly into the lipid-oil phase. The lipid oil phase comprised of 2 mM DOPC, 0.5 mM DPhPC, 0.25 mM DOPG, 0.25 mM cholesterol, dissolved in 70% hexadecane and 30% silicone oil AR20 [8]. After stabilization of the lipid monolayers, the two droplets were brought into contact to form the lipid bilayer interface (Fig. 2A). By encapsulating the pT7 LuxI, pT7 LasR, and pLas eGFP into cell 1 and pT7 LuxR, pLux LasI, and pLux eGFP into cell 2, it is possible to establish a feedforward process from cell 1 to cell 2 that subsequently drives a feedback mechanism from cell 2 to cell 1 (Fig. 2B). In cell 1, LuxI is constitutively expressed and produces 3OC6-HSL. The signaling molecule diffuses to cell 2 where it binds to the constitutively expressed transcription factor LuxR and induces expression of eGFP and LasI on two separate plasmids both containing the pLux promoter. The additional pLux eGFP plasmid was added as a marker for successful induction of the pLux promoter in cell 2. In cell 2, LasI synthesizes 3OC12-HSL which diffuses back to cell 1 to provide a feedback that induces pLas and the expression of eGFP. The dynamics of the encapsulated gene circuits within the two cells were tracked by widefield fluorescence microscopy for eGFP signal at room temperature (24-27°C). When cell 1 and 2 are in contact, eGFP expression is induced after 4.5 hours in cell 2 and after 7 hours in cell 1 (*Fig. 2C and D*). However, no eGFP expression was observed when the cells were not in contact (*see Fig. S11 in the Supplementary Materials*). This indicates that signaling between the droplets and therefore activation of the full gene circuit was only possible when the droplets were in contact with each other.

**Figure 2.**
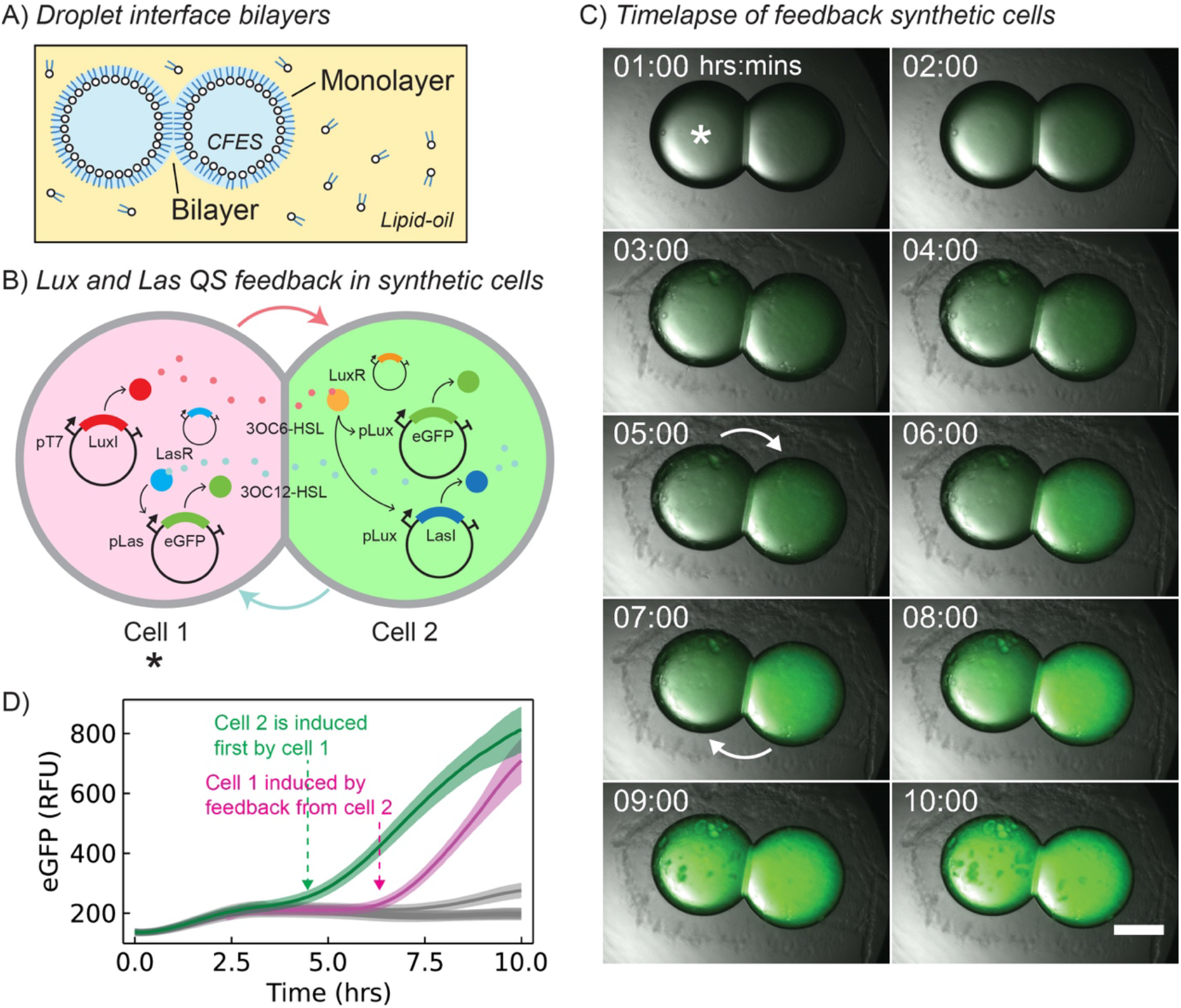
Quorum sensing feedback in droplet interface bilayers. **(A)** Illustration of droplet interface bilayers containing CFES droplets. A bilayer is formed between two water-in oil droplets stabilized by lipids when two droplets are brought into contact with one another. **(B)** Illustration of two synthetic cells encapsulating the Lux and Las QS genetic circuits to provide a feedback signaling system. Cell 1 (left, marked with an *) contains 0.5 nM pT7 LuxI, 0.5 nM pT7 LasR, and 10nM pLas eGFP. Cell 2 contains 0.5 nM LuxR, 10 nM pLux eGFP, and 0.1 nM pLux LasI. Cell 1 constitutively expresses LuxI that induces pLux eGFP and pLux LasI in cell 2. Cell 2 then induces pLux eGFP expression in cell 1 by LasI in a feedback loop. **(C)** Fluorescence widefield microscopy images of merged brightfield and eGFP fluorescence timelapse images of feedback gene circuit within DIBs over 10 hours. Cell 1 is marked with an * on the upper left image. Induction of cell 2 is marked by a white arrow at 5 hours, and the feedback induction at 7 hours. Images were obtained using a using a Zeiss Andor Axiovert 200M with a 5x/0.15 Plan-Neofluar Ph1 M27 objective. Scale bar is 500 μm. Samples were incubated at room temperature (24-27°C **(D)** Plot showing the eGFP fluorescence signal from the two cells obtained from microscopy images (B). The plot shows the initial induction of cell 2 (green) and feedback induction of cell 1 (magenta). Grey lines are the negative controls of isolated water-oil emulsion that are not in contact with each other. Shaded areas show the standard deviation of the eGFP fluorescence signal in the region of interest (ROI) in the cell used for measurement. Replicate experiments are provided in the Supplementary Materials.

The delayed induction of eGFP in cell 2 followed by cell 1 also indicates that the Lux and Las QS systems are sequentially induced by production and subsequent diffusion of the signalling molecules acyl-HSL. To verify that this was the case and that eGFP expression was not due to the diffusion of DNA, transcribed mRNA, or translated protein across the bilayer, we performed a control experiment. In the control experiment, cell 1 contained CFES in the absence of plasmid DNA and cell 2 contained CFES and a plasmid DNA for the constitutive expression of eGFP (10 nM pT7 eGFP) (*Fig. 3A and Fig. S12 in supplementary materials*). Fluorescence microscopy showed increasing fluorescence intensity in cell 2 over five hours with no increase of fluorescence intensity in cell 1 (Fig. 3B and C). These results show that the droplet interface bilayers are impermeable to DNA, mRNA, and eGFP and can therefore contain and isolate the cell free expression systems within a droplet. This indicates that expression products (DNA, mRNA, and protein) do not diffuse across the bilayer and that the presence of eGFP in the feedback gene circuit arises from the expression of lactone synthase to produce the signalling molecule acyl-HSL. Given that DNA and eGFP remain within the compartments we then wanted to confirm that the signaling molecules (acyl-HSL) could diffuse across the interface and that this mechanism was responsible for the induction of gene expression (Fig. 3D-E and Fig. S13-14 in supplementary materials). We exploited a two-cell sender and receiver system to confirm that *in situ* production of the signaling molecule acyl-HSL could diffuse across the droplet interface to induce gene expression. The sender cells contained the gene circuit to constitutively express LuxI or LasI and synthesize 3OC6-HSL or 3OC12-HSL, respectively. The receiver cells contained two plasmids, pT7 LuxR or pT7 LasR and pLux eGFP or pLas eGFP. Upon receipt of acyl-HSL from the sender cell, the transcription factor will bind and induce GFP expression. Our results show that when the sender and receiver cells were in contact with one another, positive induction and expression of eGFP was observed in both the Lux (Fig. 3Diii) and Las QS systems (Fig. 3Eiii). When the cells were not in contact, eGFP induction was only observed in the Las system (Fig. 3Div and Eiv). These results confirm that acyl-HSL produced *in situ* by constitutively expressed acyl-HSL synthases can diffuse across the bilayer to induce gene expression.

**Figure 3.**
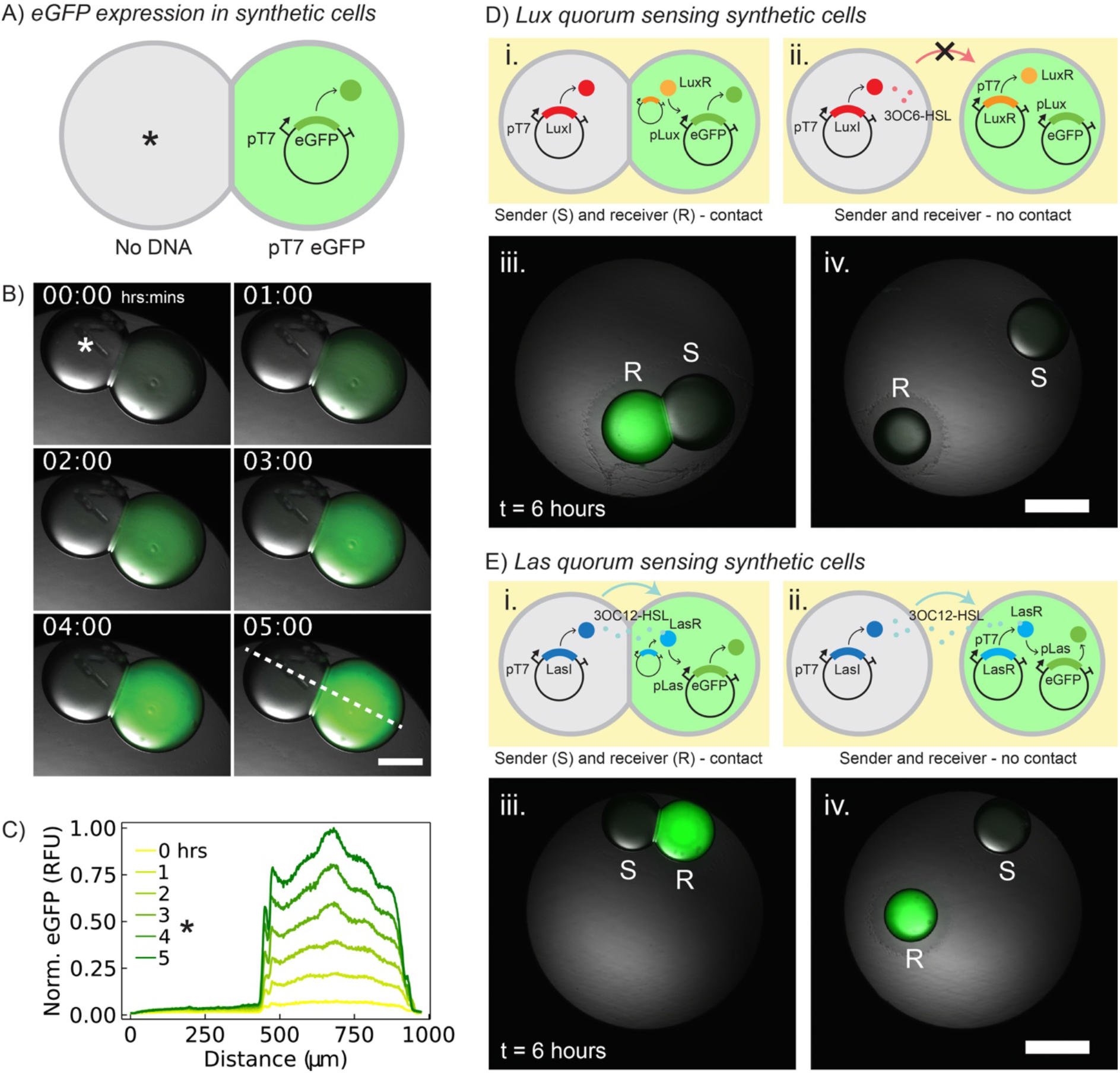
Droplet interface bilayer (DIB) synthetic cells encapsulating CFES with Lux and Las quorum sensing gene circuits. **(A)** Illustration of two DIBs encapsulating CFES without DNA plasmid (left, marked with *) and with a pT7 eGFP plasmid that constitutively expresses eGFP (right). **(B)** Fluorescence microscopy images showing merged brightfield and eGFP fluorescence timelapse images of DIB containing cell free expression system. With (10 nM) and without (*) DNA plasmid (pT7 eGFP) from 0-5 hours. Scale bar is 250 μm. **(C)** Normalized eGFP fluorescence line profiles from 0-5 hours obtained from the same pair of DIBs. The * from 0-400 μm indicates the location of the droplet without DNA plasmid. Line profile is shown on the lower right image in B. **(D)** Illustration of Lux sender (0.5 nM pT7 LuxI) and receiver (0.5 nM pT7 LuxR and 10 nM pLux eGFP) gene circuits in synthetic cells **(i)** with and **(ii)** without contact. Widefield microscopy images showing merged brightfield and eGFP fluorescence images of the Lux sender and receiver synthetic cells at 6 hours after gene expression at room temperature with **(iii)** and without contact **(iv)**. Only the Lux receiver cells in contact with the Lux sender cells are induced and express eGFP. **(E)** Illustration of Las sender (0.5 nM pT7 LasI) and receiver (0.5 nM pT7 LasR and 10 nM pLas eGFP) gene circuits in synthetic cells **(i)** with and **(ii)** without contact. Microscopy images showing merged brightfield and eGFP fluorescence images of the Las sender and receiver synthetic cells at 6 hours after gene expression. Both Las receiver cells **(iii)** with and **(iv)** without contact with the Las sender cells are induced and express eGFP. Scale bars for D and E are 500 μm. All data was acquired with a Zeiss Andor Axiovert 200M with a 5x/0.15 Plan-Neofluar Ph1 M27 objective. Samples were incubated at room temperature (24-27°C). Replicate experiments are provided in the Supplementary Materials.

Given that acyl-HSLs were previously shown to be able to partition in fluorocarbon oil (Fluorinert FC-40, Sigma Aldrich) [23,24], we hypothesize that 3OC12-HSL can more easily diffuse across the lipid monolayer interface, dissolve into the lipid-oil phase, and induce the gene expression compared to 3OC6-HSL. To test this, we characterized the partitioning of the two acyl-HSLs into the lipid-oil phase and its effect on the onset of gene expression. To do this, we added the Lux or Las receiver cell into a lipid-oil phase that had been pre-incubated with a droplet of 3OC6-HSL or 3OC12-HSL at different concentrations from 0-10000 nM (*see Fig. S15 in the Supplementary Materials*. If the acyl-HSL could partition into the lipid-oil phase and to diffuse to the receiver cell, we would expect to see eGFP induction. We observed appreciable eGFP expression in the Lux system in wells which had been preincubated with droplets containing 1000 nM of 3OC6-HSL (*see Fig. S15 BC in the Supplementary Materials*. In comparison, induction of eGFP in the Las system was observed at much lower starting concentrations of pre-incubated 3OC12-HSL droplets (10 nM). These results suggest that the lactone with the longer carbon chain is able to diffuse across the lipid monolayer interface and dissolve into the lipid-oil phase more effectively than 3OC6-HSL. Our findings suggest that it is important to note that the partitioning properties of the signaling molecules will play a significant contributing factor to the functionality of the gene circuit where the concentrations of signaling molecules are low or close to the threshold required for induction.

Taken together, our experiments describe the successful implementation of a six-plasmid gene circuit that has been separated into pairs of droplets connected by an interface bilayer. We show that the integration of gene circuits into compartments can provide additional functionality and topologies compared to a bulk reaction. Specifically, the same gene circuit in bulk runs as a sequential and linear induction, and as a feedback loop in a compartmentalized format. Our control experiments confirm that communication and signaling between the pairs of droplets are indeed driven via diffusion of the acyl-HSL signaling molecules.

### Communication in populations of synthetic cells

Finally, we wanted to extend the system from pairs of DIBs to synthetic cell communities and implement a variety of communication strategies based on different combinations of synthetic cells. Due to the difficulty in generating droplets of similar size for different cell types using a manual pipetting method, we established a new methodology to automatically dispense droplets into the lipid-oil phase and form populations of DIB synthetic cells (Fig. 4A and supplementary materials). We used the Echo 550 acoustic liquid handling machine (Echo 550, Beckman Coulter, USA) to dispense droplets between 2.5-20 nL in volume containing, the cell free expression mix, from a source well into an upside-down destination well containing the lipid-oil phase mix (Fig. 4A and supplementary materials). Between each round of droplet formation, the destination plate is flipped right side-up to allow the droplet to settle and the lipid monolayer to form and stabilize. At the end of droplet production, the 384-well destination plate was placed right-side up and tilted at an angle to allow all the droplets to come into contact and form the lipid bilayer interfaces between synthetic cells. This approach allows us to multiplex and generate custom built synthetic cell populations with a defined droplet size and population number that is specified by a transfer plan programmed to the acoustic liquid handling machine. Using this methodology, we showed two different modes of communication involving two different populations. In the first example, we took the same Lux sender (0.5 nM pT7 LuxI) and receiver (0.5 nM pT7 LuxR and 10 nM pLux eGFP) cell topology as we previously described and implemented this within a population of cells (Fig. 4B). The sender cell population was labelled with 7.2 μM purified mCherry protein for identification. After 6 hours of incubation, eGFP fluorescence was detected within all the receiver cells only (Fig. 4C), demonstrating that the quorum sensing communication functions in synthetic cell populations. Interestingly, we found that chemical induction by communication was not possible in DIBs with small 2.5 nL volume droplets (*see Fig. S16 in the Supplementary Materials*). However, control experiments showed that the constitutive gene expression of eGFP from 10 nM pT7 eGFP plasmid can take place within 2.5 nL DIBs. This could mean that, in the case of the quorum sensing systems, the concentration of signaling molecule may be diluted in the bulk lipid-oil phase to below threshold induction levels or that the higher surface-to-volume ratios in smaller droplets can somehow affect the function of the QS gene circuits.

**Figure 4.**
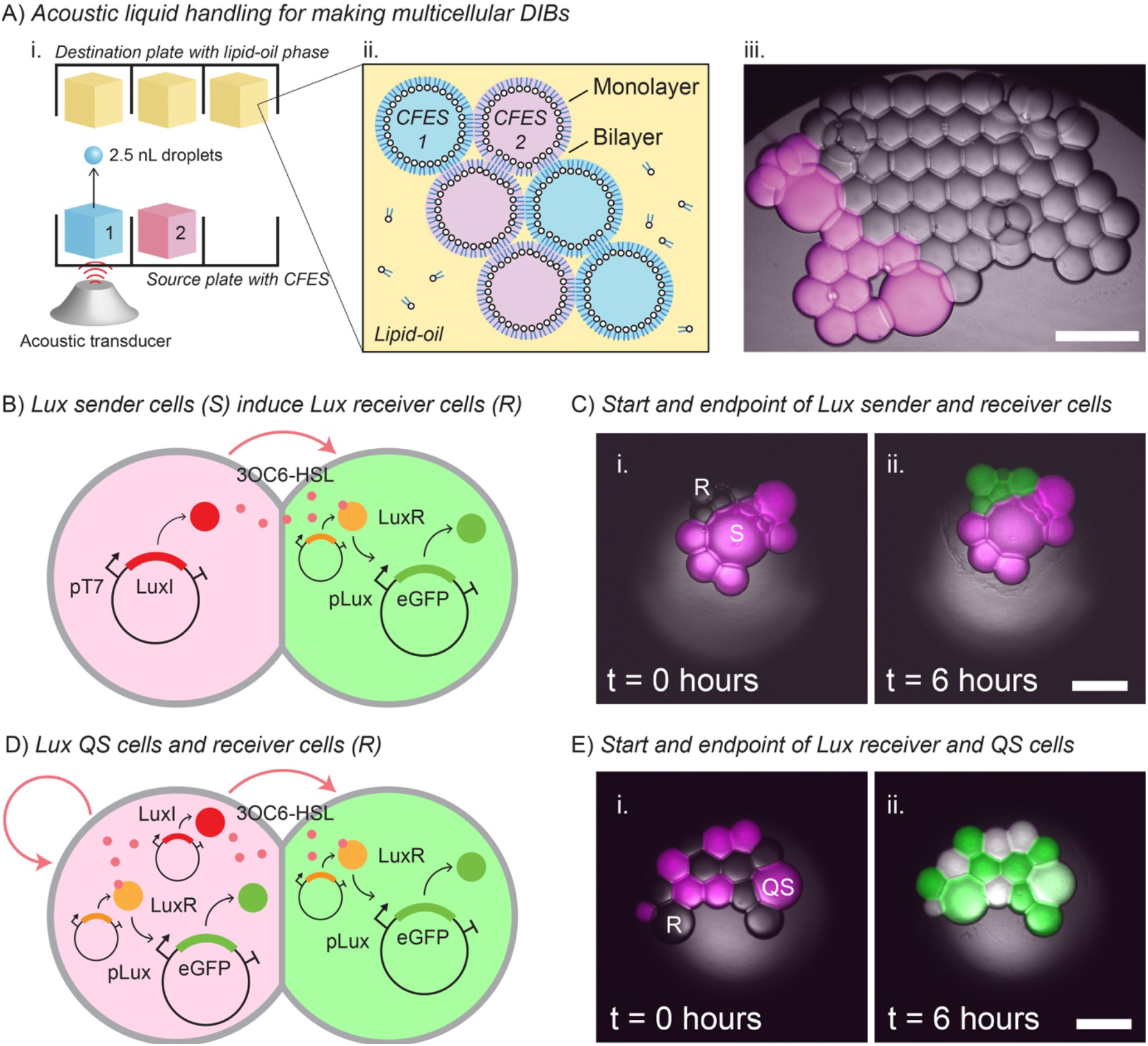
Quorum sensing communication in synthetic cell communities. **(A)** Illustration showing acoustic liquid handling for generating uniform droplets. **(i)** CFES bulk reactions are dispensed at 2.5 nL volumes into a 2 μL lipid-oil phase in a 384-well destination plate. **(ii)** Several different CFES bulk reactions can be prepared and dispensed into the same lipid-oil phase to generate heterogeneous multicellular DIBs. **(iii)** Microscopy image showing an example of a synthetic multicellular population with droplets encapsulating CFES with or without mCherry protein (magenta). **(B)** Illustration of the Lux receiver (0.5 nM pT7 LuxR and 10 nM pLux eGFP) and sender cells (0.5 nM pT7 LuxI). Sender cells were marked by adding purified mCherry protein into the CFES mix before production of the droplets. **(C)** Widefield microscopy images showing merged brightfield, mCherry, and eGFP fluorescence images of a population of Lux receiver and sender (magenta) cells at t = 0 and 6 hours. At t = 6 hours, the receiver cells show eGFP expression induced from the sender cells. **(D)** Illustration of the Lux receiver (0.5 nM pT7 LuxR and 10 nM pLux eGFP) and QS cells (0.5 nM pT7 LuxR, 10 nM pLux eGFP, and 0.5 nM pT7 LuxI). QS cells are marked by adding purified mCherry protein into the CFES mix before production of the droplets. **(E)** Widefield microscopy images showing merged brightfield, mCherry, and eGFP fluorescence images of a population of Lux receiver and QS (magenta) cells at t = 0 and 6 hours. At t = 6 hours, both receiver and QS cells show eGFP expression induced from the QS cells. Larger 20 nL droplets were created for C and E by dispensing 8 × 2.5 nL droplets in one transfer event. All scale bars are 250 μm. Data was obtained using a Zeiss Andor Axiovert 200M with a 5x/0.15 Plan-Neofluar Ph1 M27 objective (in C and E) and 10x/0.3 Plan Neo-fluar Ph1 M27 objective (in A). Samples were incubated at room temperature (24-27°C). Replicate experiments are provided in the Supplementary Materials.

In a second example, we prepared Lux receiver cells (0.5 nM pT7 LuxR and 10 nM pLux eGFP) and Lux quorum sensing cells (0.5 nM pT7 LuxI, 0.5 nM pT7 LuxR, and 10 nM pLux eGFP) (Fig. 4D). The Lux quorum sensing cells contain both the sender and receiver components of the Lux QS system. After 6 hours of incubation, GFP fluorescence was detected in both the sender and quorum sensing cells (Fig. 4E). This demonstrates the feasibility of producing heterogeneous populations of cells and coordinating their output by intra and intercellular communication. Together, our results show a novel approach to generate heterogeneous populations of cells using a commercially-available automated liquid handler where volumes and the biochemical nature of the cells can be readily tuned and customized. This opens up new possibilities to multiplex many different gene circuits into customized heterogeneous populations of synthetic cells.

## 3. Conclusion

In this study, we present our results in building bottom-up synthetic multicellular systems using quorum sensing cell-free expression systems and droplet interface bilayers. Cell-free expression systems provide a fast and open platform to test gene circuits where plasmid concentrations and cross talk between genes can be readily characterized and subsequently transferred into *in vivo* systems or synthetic compartments. Herein, we showed that cell-free gene circuits comprised of six plasmids based on the bacterial Lux and Las quorum sensing systems can be sequentially induced in bulk reactions but can also function as a feedback loop when compartmentalized into droplet interface bilayers. We show that when implementing gene circuits that it is important to consider the crosstalk between gene modules and the diffusive properties of the signaling molecules between communicating synthetic cells, and how this can affect overall circuit function. We observed different partitioning between 3OC6-HSL and 3OC12-HSL in our lipid-oil phase mixture that affects the range of communication of the Lux and Las QS systems in the synthetic cells. This is potentially beneficial for providing contact and non-contact mediated form of communication between DIB synthetic cells using the Lux and Las QS system, respectively. In addition, we provide an automated approach to build custom populations of DIB synthetic cells using a commercially available acoustic liquid handling machine. Compared to homemade 3D droplet printers [14,25], our approach can readily generate many custom synthetic cell populations into separate wells of a standard plate. On its downside, however, the spatial arrangement of these cell populations cannot be controlled. Overall, our work provides an easy assembly method to make populations of DIBs for bottom-up synthetic multicellular systems. We also showed that simple intercellular communication networks can be built using quorum sensing genetic circuits. In the future, quantification of reaction and diffusion rate parameters (*e*.*g*. calculation of Hill function parameters (*see Tables S10-11 in the Supplementary Materials*) for gene circuit design), coupled with theoretical simulations, will help the design of more complex synthetic multicellular systems using cell free systems.

## 4. Materials and methods

### Plasmid design

The quorum sensing systems from *V. fischeri* (Lux QS) and *P. aeruginosa* (Las QS) were used to create genetic circuits that allow acyl-homserine lactone (AHL) diffusion-mediated communication between synthetic cells. Separate plasmids expressing LuxI, LuxR, LasI, LasR, or eGFP proteins under transcriptional control of either T7 RNA polymerase (T7 RNAP), the Lux promoter (pLux), or the Las promoter (pLas) were constructed under the same high-copy ampicillin resistance pEXP5-NT vector backbone (Invitrogen, USA). Plasmids pEXP5-NT/6xHis eGFP, pEXP5-NT/6xHis LuxR, pEXP5-NT/6xHis LuxI, pEXP5-NT/6xHis LasR, and pEXP5-NT/6xHis LasI have a constitutive T7 RNA polymerase-mediated promoter and strong ribosomal binding site (RBS) for expression of eGFP, LuxR, LuxI, LasR, and LasI proteins, respectively. Plasmids pEXP5-NT/pLux 6xHis eGFP, pEXP5-NT/pLux 6xHis LasI, and pEXP5-NT/pLas 6xHis eGFP are controlled by the Lux or Las promoter that is induced by the LuxR:3OC6-HSL or LasR:3OC12-HSL transcription factor complex, respectively. The *luxR, luxI*, and *lasI* genes were codon optimized for *E. coli* K12 expression with a relative threshold for rare codons of 0.3 in Geneious (v11.0.2, www.geneious.com) to avoid rare codon usage. The *lasR* gene was cloned from the plasmid LasR_ADH_025 [18], which was kindly provided by Prof. Richard Murray (Caltech, USA). Plasmid construction methods and sequences are further described in the Supplementary Materials. All plasmids were sequence confirmed by Sanger sequencing and available in Addgene (www.addgene.org) with plasmid IDs 193624-193631.

### E. coli extract-based CFES

The *E. coli* BL21 (DE3) extract-based CFES was prepared using a modified protocol based on the work of Levin et al. (2019) [26]. The final 12.5 μL CFES reaction mix was composed of 5 μL extract, 1.8 μL Solution A, 1.75 μL Solution B, 0.125 μL 200 mM MgGlu, DNA plasmid templates, and water. Solution A was composed of 12.4 mM ATP (Sigma, USA), 8.7 mM GTP (Roche, Switzerland), 8.7 mM CTP (Sigma, USA), 8.7 mM UTP (Sigma, USA), 0.68 mM folinic acid (Sigma, USA), 0.176 mg/mL tRNA (Roche, Switzerland), 2.7 mM β-nicotinamide adenine dinucleotide (Sigma, USA), 1.8 mM Coenzyme A (CoA), 27.2mM oxalic acid (Roth, Germany), 6.8 mM putrescine (Sigma, USA), 10.1 mM spermidine (Sigma, USA), and 774.5mM HEPES (Roth, Germany). Solution B is composed of 14.3 mM of each of the twenty amino acids (Sigma, USA), 236.3 mM phosphoenolpyruvate (PEP), 71.6 mM magnesium glutamate (MgGlu) (Sigma, USA), and 970.8 mM potassium glutamate (KGlu) (Sigma, USA). Bulk CFES experiments with plasmid templates were prepared in a 384-well microplate, sealed with a plate film, and incubated in a TECAN Genios Pro plate reader (TECAN, Switzerland) at 27°C or 30°C for 8-12 hrs. Fluorescence levels of expressed eGFP protein were measured at emission/excitation wavelengths 485/535 nm with a gain of 25 every 5 mins. Further details of the preparation of *E. coli* extract-based CFES and bulk CFES experiments are available in the Supplementary Materials.

### CFES in DIBs

To create synthetic cells using droplet interface bilayers, CFES droplets were placed into a lipid-oil phase. The lipid-oil phase was composed of 2 mM DOPC (Avanti, USA), 0.5 mM DPhPC (Avanti, USA), 0.25 mM DOPG (Avanti, USA), and 0.25 mM cholesterol (Avanti, USA) and dissolved in 70% hexadecane (Sigma, USA) and 30% silicone oil AR20 (Sigma, USA). CFES droplets were equilibrated in the lipid-oil phase for approximately 1 minute before adding the next droplet to allow the lipid monolayer to form at the droplet interface. Droplets were made by either manually pipetting 0.05-0.1 μL droplets or dispensing individual 2.5 nL - 20 nL droplets using an Echo 550 acoustic liquid handler (Beckman Coulter, USA) into 2 μL of the lipid-oil phase in 384-well microplate. After equilibration, the plate is tilted at a 45° angle to allow the droplets to come into contact with each other and form the lipid bilayer interfaces. The surrounding wells were filled with 20 μL of water, the plate sealed with a Breathe-Easy sealing membrane (Diversified Biotech, USA), and covered with a hydration chamber to avoid droplet evaporation. The synthetic cells were incubated at room temperature (24-27°C) and imaged every 5 or 10 minutes under brightfield and widefield fluorescence microscopy using a Zeiss Andor Axiovert 200M with a 5x/0.15 Plan-Neofluar Ph1 M27 objective or 10x/0.3 Plan Neo-fluar Ph1 M27 objective. Fluorescence excitation was at 550nm through a ROX filter set (excitation bandpass 575±15 nm, beam splitter HC BS 596 nm, emission BP 641±75 nm) for mCherry or a GFP/Alexa 488/FITC filter set (excitation bandpass 449-489 nm, dichroic longpass 497 nm, emission bandpass 502-549 nm) for eGFP.

## Associated content

Supplementary material is provided in XXX. Data, code, and plasmid sequences used in this study are available in https://doi.org/10.17617/3.BCWOCD. Plasmids are also available in Addgene (<u>addgene.org</u>) with plasmid IDs 193624-193631.

## Author contributions

D.T.G., C.Z., and T-Y.D.T. designed the study. D.T.G. and S.S. performed the experiments. D.T.G. analyzed the data. D.T.G., S.S., C.Z., and T-Y.D.T. wrote the manuscript.

## Acknowledgements

We thank Javin Oza and Lei Kai for advice on making the extract-based CFES, and Aurore Dupin and Michael Booth for help in troubleshooting our DIB systems. We acknowledge financial support from the MaxSynBio Consortium which is jointly funded by the Federal Ministry of Education and Research (Germany) and the Max Planck Society. We thank the the Deutsche Forschungsgemeinschaft (DFG, German Research Foundation) under Germany’
ss Excellence Strategy – EXC-2068 – 390729961– Cluster of Excellence Physics of Life of TU Dresden and EXC-1056-Center for Advancing Electronics Dresden; Volkswagen Stiftung for generous funding. We thank the Light Microscopy Facility (LMF), the Protein Expression, Purification, and Characterization Facility (PEPC), and the Technology Development Studio (TDS) of the Max Planck Institute of Molecular Cell Biology and Genetics (MPI-CBG) for their technical support and useful discussions.

