## supplementary data for "Cell-free quorum sensing to drive feedback and communication within populations of droplet interface bilayers"

**Contents**

S1. Chemicals, materials, and equipment
S2. Strains and plasmids
S3. *E. coli* extract-based CFES
S4. Bulk CFES experiments
S5. Quorum sensing synthetic cells
S6. Echo-made synthetic cell populations

### S1. Chemicals, materials, and equipment

**Table S1.** Chemical list.

| Name | MW (g/mole) | Supplier | Catalog No. |
|---|---|---|---|
| *Amino acids* | | | |
| L-Alanine (A) | 89.09 | Sigma, USA | A7627 |
| L-Arginine (R) | 174.20 | Sigma, USA | A5006 |
| L-Asparagine (N) | 132.12 | Sigma, USA | A0884 |
| L-Aspartic acid (D) | 133.10 | Sigma, USA | A9256 |
| L-Cysteine (C) | 121.16 | Sigma, USA | W326305 |
| L-Glutamic acid (E) | 147.13 | Sigma, USA | G1251 |
| L-Glutamine (Q) | 146.14 | Sigma, USA | G3126 |
| Glycine (G) | 75.07 | Sigma, USA | G7126 |
| L-Histidine (H) | 155.15 | Sigma, USA | H8000 |
| L-Isoleucine (I) | 131.17 | Sigma, USA | I2752 |
| L-Leucine (L) | 131.17 | Sigma, USA | L8000 |
| L-Lysine (K) | 146.19 | Sigma, USA | L5501 |
| L-Methionine (M) | 149.21 | Sigma, USA | M9625 |
| L-Phenylalanine (F) | 165.19 | Sigma, USA | P2126 |
| L-Proline (P) | 115.13 | Sigma, USA | P0380 |
| L-Serine (S) | 105.09 | Sigma, USA | S4500 |
| L-Threonine (T) | 119.12 | Sigma, USA | T8625 |
| L-Tryptophan (W) | 204.23 | Sigma, USA | T0254 |
| L-Tyrosine (Y) | 181.19 | Sigma, USA | T3754 |
| L-Valine (V) | 117.15 | Sigma, USA | V0500 |
| *NTPs* | | | |
| Adenosine 5′-triphosphate disodium salt hydrate | 551.14 | Sigma, USA | A26209 |
| Cytidine 5′-triphosphate disodium salt hydrate | 527.12 | Sigma, USA | 30320 |
| Guanosine 5′-triphosphate sodium salt hydrate | 523.18 | Roche, CH | 10106399001 |
| Uridine 5′-triphosphate trisodium salt dihydrate | 586.12 | Sigma, USA | 94370 |
| *Cofactors* | | | |
| β- Nicotinamide adenine dinucleotide (NAD) | 663.43 | Sigma, USA | N1511 |
| Coenzyme A (CoA) | 767.53 | Sigma, USA | C4284 |
| Folinic acid calcium salt hydrate | 511.50 | Sigma, USA | F7878 |
| Oxalic acid | 126.07 | Roth, DE | 8879.1 |
| Phosphoenolpyruvate (PEP) | 206.1 | Roche, CH | 10108294001 |
| Putrescine | 88.15 | Sigma, USA | 51799 |
| Spermidine | 145.25 | Sigma, USA | S2626 |
| tRNA from *E. coli* MRE600 | - | Roche, CH | 10109541001 |
| *Buffers* | | | |
| Acetic acid (HOAc) | 60.05 | Merck, USA | K48001663 632 |
| Dithiothreitol (DTT) | 154.253 | Thermo, USA | R0862 |
| HEPES (N-2-Hydroxyethylpiperazine-N'-2-ethane sulphonic acid) | 238.31 | Carl Roth, DE | 9105 |
| L-Glutamic acid hemimagnesium salt tetrahydrate | 388.61 | Sigma, USA | 49605 |
| L-Glutamic acid potassium salt monohydrate | 203.23 | Sigma, USA | G1149 |
| Potassium hydroxide | 56.11 | Sigma, USA | 221473 |
| Potassium phosphate dibasic solution | 174.18 | Sigma, USA | P8584 |
| Potassium phosphate monobasic solution | 136.086 | Sigma, USA | P8709 |
| Trizma base | 121.14 | Sigma, USA | T1503 |
| *Oil phase* | | | |

| | | | |
|---|---|---|---|
| 1-Octanol | 130.23 | Sigma, USA | 297887 |
| Mineral oil | (0.84 g/mL) (1.467 n20/D) | Sigma, USA | M5904 |
| Dodecane | 170.34 | Thermo, USA | 31324 |
| Hexane | 86.18 | Sigma, USA | 139386 |
| Squalene | 410.72 | Sigma, USA | S3626 |
| Hexadecane | 226.44 | Sigma, USA | 102449651 |
| Silicone oil AR 20 | (1 g/mL) (1.441 n20/D) | Sigma, USA | 10836 |
| Hexanol | 102.17 | Sigma, USA | 471402 |
| Oleic acid | (1.459 n20/D) | Sigma, USA | 364525 |
| Undecane | (0.74 g/mL) (1.417 n20/D) | Sigma, USA | U407 |
| *Lipid phase* | | | |
| 1,2-dioleoyl-sn-glycero-3-phosphocholine (DOPC) | 786.113 | Avanti, USA | 850375C |
| 1,2-diphytanoyl-sn-glycero-3-phosphocholine (DPhPC) | 846.252 | Avanti, USA | 850356C |
| 1,2-dioleoyl-sn-glycero-3-phospho-(1'-rac-glycerol) (sodium salt) (DOPG) | 797.026 | Avanti, USA | 840475 |
| Cholesterol | 386.65 | Sigma, USA | C3045 |
| *Others* | | | |
| Twinsil Speed silicone | - | Picodent, DE | 1300 1002 |

**Table S2.** Kit list.

| Name | Supplier | Catalog No. |
|---|---|---|
| NEBuilder HiFi DNA Assembly Kit | NEB, USA | E2621 |
| QIAprep Spin Miniprep Kit | QIAGEN, DE | 27104 |
| QIAGEN Plasmid Maxi Kit | QIAGEN, DE | 12162 |
| QIAquick PCR Purification Kit | QIAGEN, DE | 28104 |
| myTXTL Sigma 70 Master Mix Kit | Daicel Arbor Biosciences, USA | 507024 |

**Table S3.** Consumables list.

| Name | Supplier | Catalog No. |
|---|---|---|
| 384-well plates, Lobase, Black | Greiner Bio-One, AT | 788096 |
| Adhesive PCR Plate Seals | Thermo, USA | AB0558 |
| Breathe-Easy sealing membrane | Diversified Biotech, USA | Z380059 |
| Spectra/Por 2 Dialysis Membrane (12-14 kD) | Repligen, USA | 132680T |
| Echo Qualified 384-Well Low Dead Volume (LDV) Source Microplate | Beckman Coulter, USA | 001-12782 |

**Table S4.** Equipment list.

| Name | Supplier | Catalog No. |
|---|---|---|
| 5x/0.15 Plan-Neofluar Ph1 M27 objective | Zeiss, DE | 420331-9911-000 |
| 10x/0.3 Plan Neo-fluar Ph1 M27 objective | Zeiss, DE | 420341-9911-000 |
| Andor Axiovert 200M | Zeiss, DE | - |
| Avanti Centrifuge J26-XP | Beckman Coulter, USA | - |
| JLA-8.1000 rotor | Beckman Coulter, USA | 363688 |
| NanoDrop 2000 | Thermo, USA | ND-2000 |
| pE-4000 illumination system | CoolLED, USA | - |
| Sonorex sonicator bath | Bandelin, DE | - |
| Branson Digital Sonifier 450-D (with 418-A probe) | Thermo, USA | - |
| Genios Pro | TECAN, CH | - |
| Echo 550 acoustic liquid handler | Beckman Coulter, USA | - |

### S2. Strains and plasmids

The *E. coli* bacterial strains and plasmids used in this study are listed in Table S5-6. The plasmid pEXP5-NT/6xHis eGFP was provided by JLR Anderson, University of Bristol [1] and used as a positive control for T7 RNA polymerase (RNAP)-mediated expression of eGFP. The plasmid pTXTL-P70a-deGFP, provided in the myTXTL Sigma 70 Master Mix Kit (Daicel Arbor Biosciences, USA), was used as a positive control for native or core RNAP and $\sigma^{70}$-mediated expression of deGFP. The plasmid LasR_ADH_025, provided by RM Murray and AD Halleran [2], was used as a PCR template to obtain the *lasR* gene for cloning. Codon-optimized sequences for *luxR*, *luxI*, and *lasI* genes were designed in Geneious 11.0.2 (www.geneious.com) using the standard *E. coli* K12 genetic code with a rare codon threshold of 0.3 and synthesized as double-stranded DNA gBlocks (IDT, USA). Plasmids constructed in this study used the pEXP5-NT vector backbone of the pEXP5-NT/6xHis eGFP plasmid. Vector and insert parts were made by PCR amplification and overlap extension (NEB Phusion HF). Plasmid assembly of the vector and insert parts was done by Gibson assembly using the NEBuilder HiFi DNA Assembly Kit (NEB, USA) or restriction digest and ligation. All plasmids were prepared and purified by ethanol precipitation from *E. coli* DH5*α* cultures using the QIAGEN Plasmid Maxi Kit (QIAGEN, Germany) and measured by NanoDrop 2000 (Thermo, USA). All PCR, purification, and assembly methods were performed using the manufacturer's standard protocols. All primers, ultramers, and gBlocks were synthesized by Integrated DNA Technologies (IDT, www.idtdna.com). Plasmid assembly was confirmed by Sanger sequencing (GENEWIZ, www.genewiz.com) and deposited in Addgene (addgene.org) with plasmid IDs 193624-193631. Plasmid sequences in GenBank format are provided in the supplementary files.

**Table S5.** Strains used in this study.

| Strain name | Description | Source |
|---|---|---|
| *E. coli* DH5α | High efficiency competent cells for transformation. | NEB (C2987) |
| *E. coli* BL21 (DE3) | Competent cells for T7 RNAP-mediated expression. | NEB (C2527) |

**Table S6.** Plasmids used in this study.

| Plasmid name | Simple name | Description | Source |
|---|---|---|---|
| pEXP5-NT/6xHis eGFP | pT7 eGFP | Constitutive T7 RNAP-mediated expression of 6xHis eGFP. | [1] |
| pTXTL-P70a-deGFP | p70a deGFP | Constitutive core RNAP and $\sigma^{70}$-mediated expression of 6xHis eGFP. | Daicel Arbor Biosciences, USA |
| LasR_ADH_025 | - | Used for PCR extension of *lasR* gene. | [3] |
| pEXP5-NT/pT7 LuxR | pT7 LuxR | Constitutive T7 RNAP-mediated expression of 6xHis LuxR. Addgene ID 193625. | This study. |
| pEXP5-NT/pT7 LuxI | pT7 LuxI | Constitutive T7 RNAP-mediated expression of 6xHis LuxI. Addgene ID 193626. | This study. |
| pEXP5-NT/pLux eGFP | pLux eGFP | Expression of 6xHis eGFP driven by the Lux promoter. Addgene ID 193624. | This study. |
| pEXP5-NT/pT7 LasR | pT7 LasR | Constitutive T7 RNAP-mediated expression of 6xHis LasR. Addgene ID 193629. | This study. |
| pEXP5-NT/pT7 LasI | pT7 LasI | Constitutive T7 RNAP-mediated expression of 6xHis LasI. Addgene ID 193631. | This study. |
| pEXP5-NT/pLas eGFP | pLas eGFP | Expression of 6xHis eGFP driven by the Las promoter. Addgene ID 193630. | This study. |
| pEXP5-NT/pLux LasI | pLux LasI | Expression of 6xHis LasI driven by the Lux promoter. Addgene ID 193628. | This study. |

### S3. *E. coli* extract-based CFES

The *E. coli* extract-based CFES is prepared using a modified protocol based on the work of Levin *et al.* (2019) [4]. The final CFES mix is composed of the extract, Solution A, Solution B, and the DNA template for gene expression. Solution A contains the NTPs, tRNAs, and cofactors. Solution B contains the amino acids, energy regeneration system, and glutamate salts. The main differences between this protocol and of Levin *et al.* (2019) include the following: (i) no glucose supplement in the 2xYTP production culture media to improve gene expression activity of the extract [5], (ii) no IPTG induction of production culture (we rely on the leaky expression of T7 RNAP expression in *E. coli* BL21 (DE3)), (iii) including a post-processing dialysis step in S30B buffer after the run-off reaction to improve transcriptional activity of native gene expression [5], and (iv) no ammonium glutamate in Solution B, a lower concentration of tRNA, and other minor differences in the final concentrations of Solutions A and B, mostly due to convenience and availability of chemicals/materials in the lab.

*E. coli extract*

Prepare the following materials beforehand: (i) 200 mL LB media, (ii) 1 L 2xYTP media (5 g NaCl, 10 g yeast extract, 16 g tryptone, 40 mL 1 M potassium phosphate dibasic solution, 22 mL 1 M potassium phosphate monobasic solution, fill with water to 1 L), (iii) 500 mL S30A buffer (14 mM MgGlu, 60 mM KGlu, 50 mM Tris, and 2 mM DTT, titrated with ~1 mL glacial acetic acid to pH 7.7), (iv) 1L S30B buffer (14 mM MgGlu, 60 mM KGlu, and 2 mM DTT, titrated with ~2 mL 2 M Tris to pH 8.2), and (v) 1 mL 1 M DTT. Sterilize S30A and S30B solutions by autoclave and only add DTT before use. For *E. coli* extract preparation, all the following steps are done on ice or 4°C unless otherwise stated.

1) Starter culture: Inoculate *E. coli* BL21 (DE3) in 200 mL LB media and incubate overnight at 37°C with 180 rpm shaking.
2) Production culture: Use the starter culture to inoculate two production cultures of pre-warmed 500 mL 2xYTP media each at a starting OD600 of 0.05. Incubate at 37°C with 180 rpm shaking until an OD600 of 1.6 (approx. 3-5 hours).
3) Pellet the cells by centrifugation for 10 mins at 5000 xg using an Avanti Centrifuge J26-XP with a JLA-8.1000 rotor (Beckman Coulter, USA).
4) Scoop out the pellet to a pre-weighed 50 mL tube and wash three times with 30 mL S30A buffer.
5) Weigh the washed pellet ~2.5g of wet pellet/500 mL culture), flash freeze with liquid nitrogen, and store at -80°C overnight.
6) The next day, add 1 mL S30A buffer per 1 g wet pellet, thaw, and resuspend by vortexing.
7) Transfer resuspended cells into 1.5 mL aliquots in 2 mL microcentrifuge tubes.
8) Sonicate cells in an ice-water bath to prevent overheating of the sample. Sonicate by 10 cycles of 10 s pulse and 30 s rest at 25% amplitude (Branson Digital Sonifier Model 450-D with a 418-A probe). The resulting lysate should turn a darker brown compared with the pre-sonicated resuspension.
9) Immediately after sonication, add 2 µL of 1M DTT into the 2 mL microcentrifuge tube.
10) Centrifuge the sonicated lysates at 18,000 xg for 10 mins.
11) Run-off reaction: Collect and pool the clear supernatant into a 10mL tube and incubate at 37°C and 250 rpm with the cap open for 1 hour.
12) Clarify the lysate by centrifugation at 10,000 xg for 10 mins to remove precipitates.
13) Dialyze the clarified lysate in S30B buffer for 3 hours at 4°C using a Spectra/Por 2 Dialysis Membrane with a 12-14 kDa molecular weight cut-off (Repligen, USA).
14) Clarify the dialyzed lysate by centrifugation at 10,000 xg for 10 mins.
15) Aliquot the lysates in 50 µL volumes and flash freeze in liquid nitrogen. Store at -80°C until use.

*Solution A*

The stock and final concentrations of the components used for Solution A are listed in Table S7. All stock solutions were prepared by dissolving in water and the HEPES buffer was adjusted to pH 7 using KOH. 50 µL aliquots of Solution A were flash frozen with liquid nitrogen and stored at -80°C until use.

**Table S7.** Final mix for Solution A.

| Component | MW (g/mole) | Stock (mM) | Final (mM) |
|---|---|---|---|
| ATP | 507.18 | 100 | 12.4 |
| GTP | 523.18 | 100 | 8.7 |
| CTP | 483.156 | 100 | 8.7 |
| UTP | 484.1411 | 100 | 8.7 |
| Folinic acid | 473.44 | 21 | 0.68 |
| tRNA | - | 50 mg/mL | 0.176 mg/mL |
| NAD | 663.43 | 100 | 2.7 |
| CoA | 767.53 | 50 | 1.8 |

| | | | |
|---|---|---|---|
| Oxalic acid | 184.23 | 1000 | 27.2 |
| Putrescine | 88.15 | 200 | 6.8 |
| Spermidine | 145.25 | 250 | 10.1 |
| HEPES (pH 7) | 238.3012 | 2000 | 774.5 |

*Solution B*

The stock and final concentrations of the components of Solution B are listed in Table S8. The amino acid mix contains 50 mM of *each* amino acid and is incubated at 37°C with shaking to help dissolve the powder. Note that the amino acids may not dissolve completely in solution. The amino acid mix is prepared in a large volume of 10 mL to ensure that sufficient material could be weighed out. The amino acid solution should be fully mixed before adding into the final Solution B mixture. The stock of magnesium glutamate and potassium glutamate were prepared as one solution. The PEP stock solution was adjusted to pH 7 using 10 M KOH solution. The amounts of the final Solution B mix are given to make 2.1 mL of Solution B or approximately 1000 reactions. 50 µL aliquots of Solution B were flash frozen with liquid nitrogen and stored at -80°C until use.

**Table S8.** Final mix for Solution B.

| Component | MW (g/mole) | Stock (mM) | Final (mM) |
|---|---|---|---|
| Amino acid mix | - | 50 | 14.3 |
| Magnesium glutamate | 388.61 | 150 | 71 |
| Potassium glutamate | 203.23 | 2000 | 930 |
| PEP (pH 7) | 206.1 | 1000 | 238 |

*Running the CFES reaction mix*

The final *E. coli* extract-based CFES master mix is prepared according to Table S9 and incubated at 25-37°C for 5-10 hours. The amount of magnesium glutamate supplemented to the final CFES mix varies from batch to batch (0-20 mM) and was optimized to be 2mM. To prepare the CFES mix, the DNA + water components of each sample were first pre-mixed, the remainder of the master mix was then added to avoid long waiting times between starting the CFE reactions. All bulk and encapsulated experiments in this study follow the CFES mix in Table S9.

**Table S9.** A standard *E. coli* extract-based CFES master mix

| Component | µL |
|---|---|
| Solution A | 1.8 |
| Solution B | 1.75 |
| Extract | 5 |
| MgGlu (200 mM) | (0.125) |
| Plasmid DNA | x |
| Water | Fill to 12.5 |

### S4. Bulk CFES experiments

For all bulk CFES experiments, a CFES mix was prepared according to Table S7 to a total volume of 12.5 μL in a 384-well microplate, sealed with a clear PCR plate film. The well plate was incubated in a TECAN Genios Pro plate reader at 27°C or 30°C for 4-12 hrs. Fluorescence levels of expressed eGFP protein were measured at emission/excitation wavelengths 485/535 nm with a gain of 25 every 5 mins.

*Batch controls and calibration*

To convert fluorescence levels of eGFP to concentration units, a series dilution of purified eGFP [6] with known concentration was prepared in the CFES master mix to obtain a calibration curve from the plate reader (Fig. S1A). This allowed us to estimate the concentration of expressed eGFP in all our bulk experiments. To test our *E. coli* extract-based CFES, two positive control plasmids that constitutively expresses eGFP protein using T7 RNAP (pEXP5-NT/eGFP [1]) or expresses deGFP using the native *E. coli* RNAP (p70a-deGFP) is used to check for active expression (Fig. S1B). During the course of this work, we prepared and used two batches of CFES and compared their expression yields (Fig. S2C). Although we used two different batches throughout the study, all controls for each set of experiments were performed within the same batch.

*Lux and Las quorum sensing systems*

Plasmid concentrations of the Lux and Las quorum sensing systems were chosen from initial test experiments by titrating individual plasmid concentrations. The concentration of the plasmids in the reaction mix were selected based on high eGFP expression (Figure S2). Concentrations were set at 10 nM pLux eGFP, 0.5 nM pT7 LuxR, and 0.5 nM pT7 LuxI for the Lux QS system and 10 nM pLas eGFP, 0.5 nM pT7 LasR, and 0.5 nM pT7 LasI for the Las QS system unless otherwise stated. To confirm whether the pLux eGFP and pLas eGFP plasmids can be induced by their respective transcription factors and acyl-HSLs or acyl-HSL synthases. 3OC6-HSL/3OC12-HSL (0-1000 nM) and pT7 LuxI/pT7 LasI (0-1 nM) on the Lux and Las quorum sensing systems were titrated into a cell free expression mix containing 0.5 nM pT7 LuxR and 10 nM pLux eGFP (Fig. S3-4, endpoint data is shown in Fig. 1C-D in the main text). Titration experiments of 3OC6-HSL/3OC12-HSL and pT7 LuxI/pT7 LasI on the Lux and Las quorum sensing systems were additionally used to estimate the parameters of the Hill function $[Y] = k\,[X]^n/(K_d + [X]^n)$ (Tables S10-11). However, only the titration of the Lux quorum sensing system with pT7 LuxI resulted in identifiable parameters as indicated by its confidence intervals.

*Cross-talk of Lux and Las QS systems*

It was previously reported that the Lux and Las quorum sensing systems show cross-talk between each other under cell free conditions [2]. We validated this with our gene circuits and *E. coli* extract-based CFES. Endpoint eGFP expression of the Lux and Las quorum sensing gene circuits were obtained after induction by 3OC6-HSL or 3OC12-HSL (1000 nM) (Fig. S6A and C) and pT7 LuxI or pT7 LasI plasmids (0.5 nM) (Figs. S6B and D). Our results show cross-talk of the Lux QS with 3OC12-HSL but not the Las QS with 3OC6-HSL.

*Leaky expression of pLux LasI plasmid for series induction*

For the series induction of Lux and Las QS systems, we also ensured that the concentration of pLux LasI plasmid used does not result in leaky expression of LasI. Different concentrations of the pLux LasI plasmid was tested to see which concentrations can lead to premature induction of the Las quorum sensing system. From 0.1-5 nM of pLux LasI plasmid, only the concentration 0.1 nM did not lead to leaky induction of the Las quorum sensing system (Fig. S7).

*Room temperature controls*

Bulk experiments were repeated to confirm that the qualitative behavior of the cross-talk of the quorum sensing gene circuits and Lux-Las series induction remained the same at lower temperatures (approximately 24-27°C) as encapsulated experiments were undertaken at room temperature (Fig. S8).

*Oil compatibility with CFES*

The compatibility of different oils (mineral oil, hexane, squalene, octanol, hexanol, oleic acid) that can be used to prepare DIBS was tested against the *E. coli* extract-based CFES. To do this, 5 μL of oil was overlayed on top of a standard bulk CFES reaction expressing eGFP from 5 nM pT7 eGFP plasmid. Endpoint expression (t = 4 hours) of the different oils used show varying degrees of reduction of eGFP expression (Fig. S9).

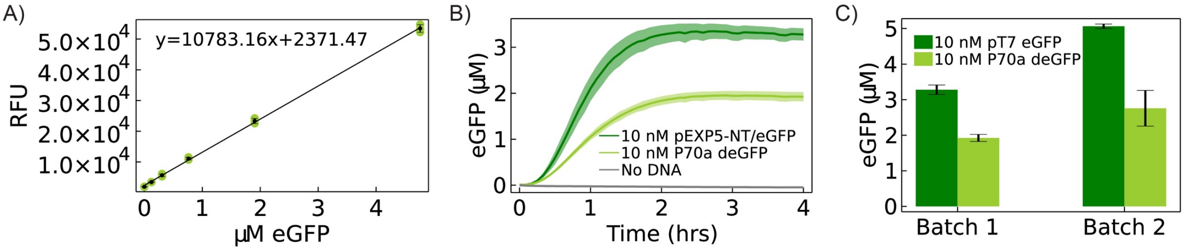


**Figure S1.** (A) Calibration of eGFP protein (µM) vs. RFU in *E. coli* extract-based CFES. Green data points show triplicate samples. Black dots and error bars are the mean and standard deviation values. The solid black line is the linear fit of the calibration curve y = 10783.16 x + 2371.47. (B) eGFP protein levels over time expressed from 10 nM pT7 eGFP, 10 nM P70a deGFP, or no plasmid DNA in Batch 1 of the *E. coli* extract-based CFES incubated at 30°C. (C) Comparison of endpoint eGFP expression (t = 4 hours) from 10 nM pT7 eGFP and 10 nM P70a deGFP plasmids between Batch 1 and 2 of the *E. coli* extract-based CFES.

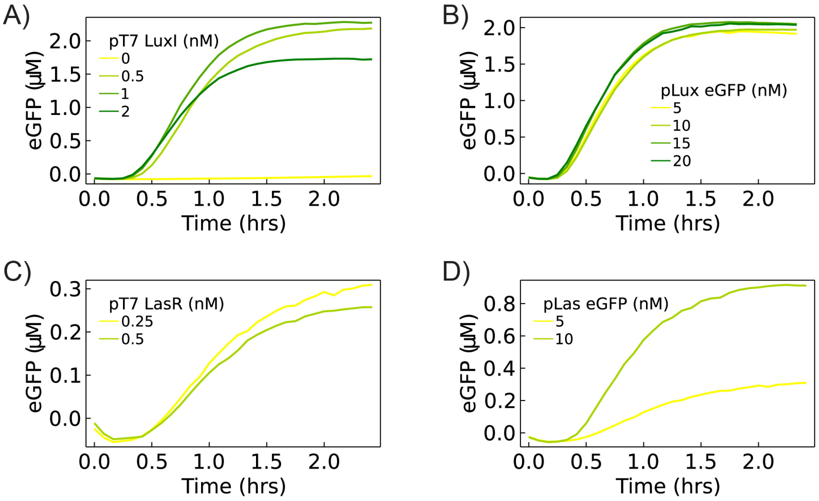


**Figure S2.** Optimizing plasmid concentrations for the Lux and Las quorum sensing system. (A) Titration of pT7 LuxI plasmid from 0-2 nM with 10 nM pLux eGFP and 0.5 nM pT7 LuxR. (B) Titration of pLux eGFP plasmid from 5-20 nM with 2 nM pT7 LuxR and 2 nM pT7 LuxI. (C) 0.25 or 0.5 nM pT7 LasR plasmid with 5 nM pLas eGFP and 100 nM 3OC12-HSL. (D) 5 or 10 nM pLas eGFP plasmid with 0.25 nM pT7 LasR and 100 nM 3OC12-HSL.

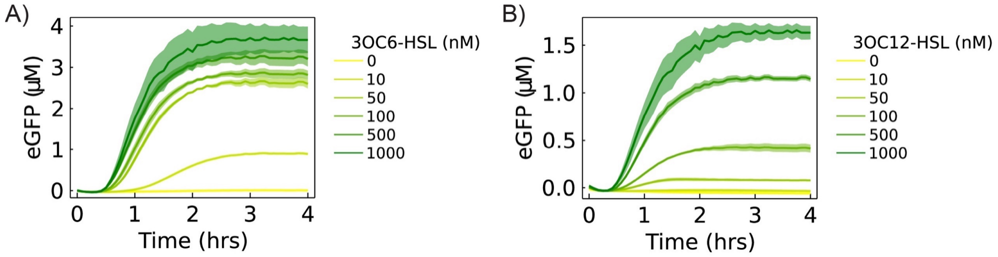


**Figure S3.** eGFP expression levels over time with (A) 0-1000 nM 3OC6-HSL into the Lux quorum sensing system (0.5 nM pT7 LuxR and 10 nM pLux eGFP) and (B) 0-1000 nM 3OC12-HSL into the Las quorum sensing system (0.5 nM pT7 LasR and 10 nM pLas eGFP) in *E. coli* extract-based CFES. Solid lines and shaded areas indicate the mean and standard deviation from triplicate experiments.

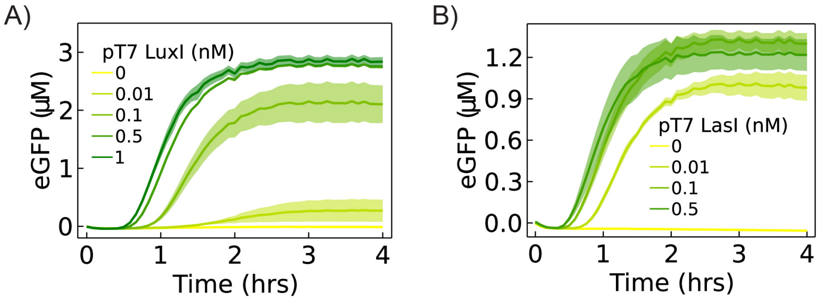


**Figure S4.** (A) eGFP expression levels over time of (A) 0-1 nM pT7 LuxI plasmid into the Lux quorum sensing system (0.5 nM pT7 LuxR and 10 nM pLux eGFP) and (B) 0-0.5 nM pT7 LasI plasmid into the Las quorum sensing system (0.5 nM pT7 LasR and 10 nM pLas eGFP) in *E. coli* extract-based CFES. Solid lines and shaded areas indicate the mean and standard deviation from triplicate experiments.

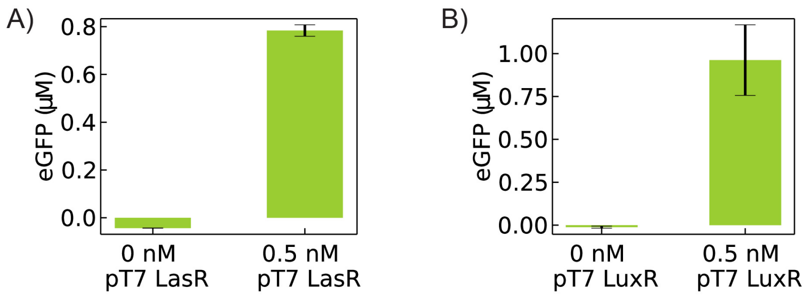


**Figure S5.** Endpoint (t = 4 hours) eGFP levels of the (A) Lux quorum sensing system (10 nM pLux eGFP and 0.5 nM pT7 LuxI) with and without 0.5 nM of pT7 LuxR plasmid and (B) Las quorum sensing system (10 nM pLas eGFP and 0.5 nM pT7 LasI) with and without 0.5 nM of pT7 LasR plasmid. Error bars indicate the standard deviation from triplicate samples.

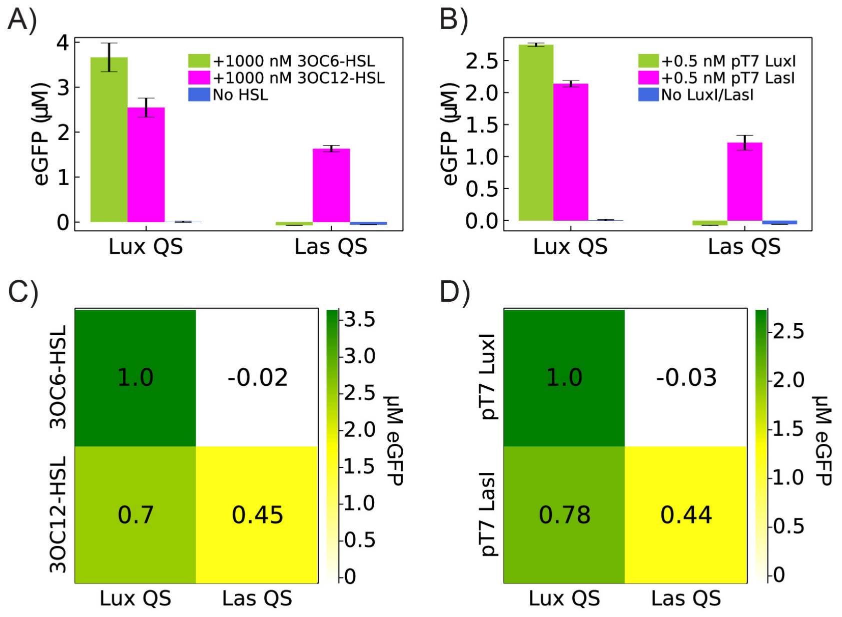


**Figure S6.** Cross-talk between Lux and Las quorum sensing systems. Endpoint (t = 4 hours) concentrations of expressed eGFP to show cross-talk levels of Lux (0.5 nM pT7 LuxR and 10 nM pLux eGFP) and Las quorum sensing systems (0.5 nM pT7 LasR and 10 nM pLas eGFP) against (A) 1000 nM of 3OC6-HSL or 3OC12-HSL or (B) 0.5 nM pT7 LuxI or pT7 LasI plasmids. Heatmap of the same data showing the cross-talk of the Lux and Las quorum sensing systems against (C) 3OC6-HSL or 3OC12-HSL and (D) 0.5 nM pT7 LuxI or 0.5 nM pT7 LasI plasmids. Color of each square indicates the concentration of eGFP expressed at endpoint (t = 4 hours) as noted in the color bar on the right. Values written within each square is the normalized eGFP concentration with respect to the highest value given by the expression of the Lux system induced by 3OC6-HSL for A or pT7 LuxI for B.

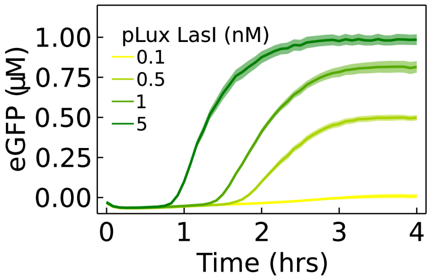


**Figure S7.** Leaky expression and induction of the pLux LasI plasmid from 0.1-5 nM against the Las quorum sensing system (0.5 nM p7 LasR and 10 nM pLas eGFP).

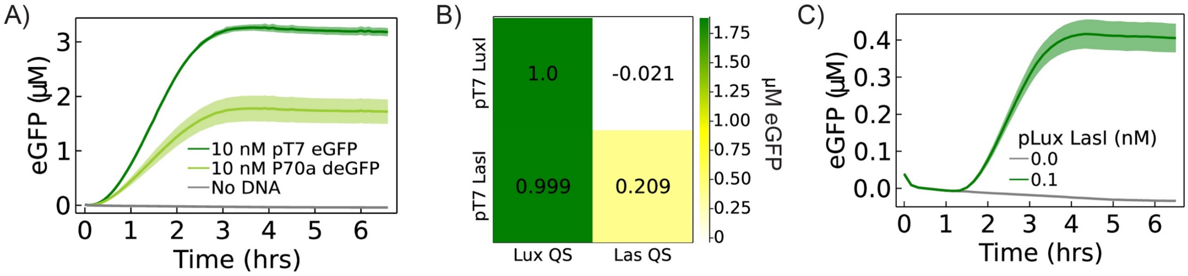


**Figure S8.** Repeat of bulk CFES experiments set at a lower temperature (27°C) for (A) constitutive expression from 10 nM pT7 eGFP and 10 nM P70a deGFP, (B) cross-talk of the Lux and Las quorum sensing systems, and (C) Lux-Las series induction showing the same qualitative behavior of the gene circuits as the same experiments undertaken at 30 ºC.

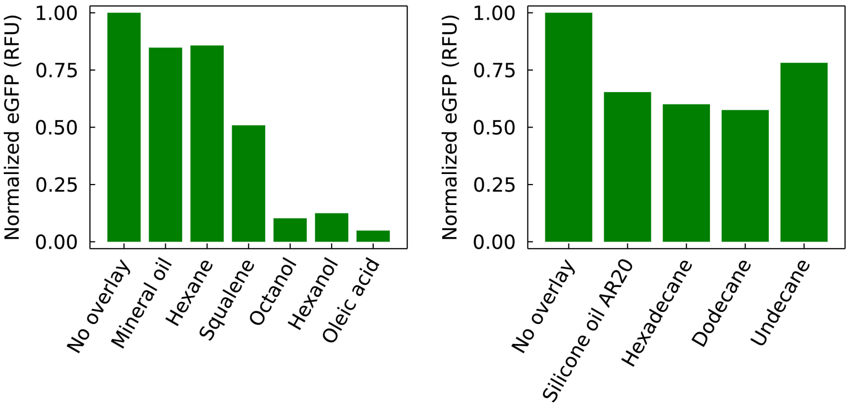


**Figure S9.** Normalized eGFP endpoint fluorescence (t = 4 hours) for different oils overlaid on E. coli extract-based CFES reactions with 5 nM pT7 eGFP plasmid.

**Table S10.** Hill parameter fit and confidence intervals for Lux quorum sensing system.

| Parameter | pT7 LuxI | | | 3OC6-HSL | | |
|---|---|---|---|---|---|---|
| | LSE | 95% CI | | LSE | 95% CI | |
| | | lower bound | upper bound | | lower bound | upper bound |
| k | 2.864 | 2.801 | 2.928 | 3.498 | 2.929 | 4.068 |
| $K_d$ | 0.014 | 0.008 | 0.020 | 39.362 | -53.873 | 132.597 |
| n | 1.419 | 1.276 | 1.561 | 1.160 | 0.405 | 1.914 |

**Table S11.** Hill parameter fit and confidence intervals for Las quorum sensing system.

| Parameter | pT7 LasI | | | 3OC12-HSL | | |
|---|---|---|---|---|---|---|
| | LSE | 95% CI | | LSE | 95% CI | |
| | | lower bound | upper bound | | lower bound | upper bound |
| k | 1.259 | -26.581 | 29.098 | 2.377 | -0.115 | 4.869 |
| $K_d$ | 0.000 | -5.379 | 5.379 | 1148.400 | -3195.033 | 5491.832 |
| n | 2.587 | -1215949.751 | 1215954.924 | 1.130 | 0.203 | 2.057 |

### S5. Quorum sensing synthetic cells

Synthetic cell pairs were made by incorporating the *E. coli* extract-based CFES into droplet interface bilayers (DIBs) by manually pipetting two droplets each with volumes of approximately 0.05-0.1 µL into 10 µL of the lipid-oil phase in a 384-well microplate. The lipid-oil phase was composed of 2 mM DOPC, 0.5 mM DPhPC, 0.25 mM DOPG, and 0.25 mM cholesterol and dissolved in 70% hexadecane and 30% silicone oil AR20. The formulation was based on the work of Dupin and Simmel (2019) [7], but with the ratio of hexadecane and silicone oil AR20 modified. To prepare the lipid-oil phase, the appropriate amounts of DOPC, DPhPC, DOPG, and cholesterol in chloroform stock solutions were mixed together in a glass test tube, dried under flowing nitrogen gas for 5 minutes and dried under vacuum for 30 minutes. The dry lipid film was resuspended in 1 mL of 70:30 hexadecane:silicone oil AR20, and incubated at 37 °C in a sonication bath (Sonorex sonicator bath) for 30 minutes. The CFES droplets were equilibrated in the lipid-oil phase for approximately 5 minutes before letting the two droplets come into contact to allow the lipid monolayer to form at the droplet interface. Note that a longer equilibration time was used compared to the smaller Echo-made droplets as larger droplets were observed to be less stable during DIBs production. After equilibration, the plate is tilted at a 45° angle to allow the droplets to come into contact with each other and form the lipid bilayer interfaces. The surrounding wells were filled with 20 µL of water, the plate is sealed with a Breathe-Easy sealing membrane, and covered with a hydration chamber to avoid droplet evaporation (Fig. S10). The synthetic cells were imaged under brightfield and widefield fluorescence microscopy using a Zeiss Andor Axiovert 200M with a 5x/0.15 Plan-Neofluar Ph1 M27 or 10x/0.3 Plan Neo-fluar Ph1 M27 objective. Fluorescence excitation was at 550 nm through a ROX filter set (excitation bandpass 575±15 nm, beam splitter HC BS 596 nm, emission BP 641±75 nm) for mCherry or a GFP/Alexa 488/FITC filter set (excitation bandpass 449-489 nm, dichroic longpass 497 nm, emission bandpass 502-549 nm) for eGFP. All synthetic cell experiments were done at room temperature between 24-27 °C and imaged at 10-minute intervals over 10 hours. Note that initial timepoints are delayed as sample preparation of the synthetic cells can typically take 30 minutes to 1 hour.

*Feedback quorum sensing cells*

Feedback synthetic cells were made by encapsulating the Lux-Las series induction plasmids in separate DIB compartments as shown in Fig. 2B in the main text. We avoid premature induction of the feedback because LuxI cannot induce the Las QS system by cross-talk (Fig. S6). Control experiments with Cell 1 and Cell 2 in isolation or within the same well but not in contact were undertaken to show that there is no self-induction or leaky induction and that both cells in contact are required for feedback induction (Fig. S11).

*eGFP expression*

To test the permeability of the gene expression products (DNA, mRNA, protein) between connected droplet interface bilayers, two droplets with CFES with and without 10 nM of pT7 eGFP plasmid were placed in contact with each other. If DNA, mRNA, or protein are permeable to the lipid bilayer membrane between the two cells, then both cells should exhibit an increase in eGFP fluorescence. However, eGFP signal only increases in the cell with the pT7 eGFP plasmid, which indicates that the gene expression products remain within its compartment (Fig. S12).

*Lux and Las quorum sensing cells*

The Lux and Las quorum sensing gene circuits were tested by encapsulating the sender and receiver components of the quorum sensing system into separate DIB compartments. The sender cell contains the plasmid to express the HSL synthase (0.5 nM pT7 LuxI or 0.5 nM pT7 LasI) and the receiver cell contains the inducible components (0.5 nM pT7 LuxR and 10 nM pLux eGFP or 0.5 nM pT7 LasR and 10 nM pLas eGFP). In isolation, the receiver cells are not induced and do not express eGFP. The Lux receiver cell is only induced when in direct contact with the Lux sender cell (Fig. S13). In contrast, the Las receiver cell is induced even if the Las sender cell is not in direct contact (Fig. S14). We hypothesized that the 3OC12-HSL from the Las sender cell partitions into the oil phase and then into the Las receiver cell to induce eGFP expression. To test this, we preincubated 0.25 µL droplets of different concentrations (0-1000 nM) of 3OC12-HSL or 3OC6-HSL into 10 µL of the lipid-oil phase for 2 hours. 7.5 µL of the lipid-oil phase without the HSL droplet was then transferred to an empty clean well to which a receiver cell was added. The receiver cell will be induced if enough HSL can partition into the lipid-oil phase and into the receiver cell. Induction of the gene was detected by eGFP fluorescence. It was found that the Las receiver cell was induced at lower concentrations of preincubated 3OC12-HSL droplets compared to the Lux receiver cell with preincubated 3OC6-HSL droplets (Fig. S15). This was unexpected as the Lux QS system was found to be induced at lower concentrations of 3OC6-HSL compared to the Las QS system with 3OC12-HSL in bulk experiments (Figs. 1C-D in the main text).

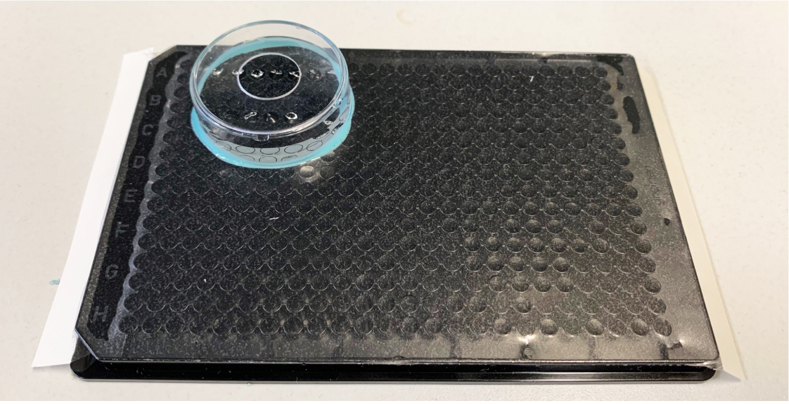

**Figure S10.** Hydration chamber setup of synthetic cell experiments. After preparing the samples, the surrounding wells were filled with 20 μL of water, the plate is sealed with a Breathe-Easy sealing membrane, and covered with a hydration chamber. The hydration chamber is composed of an inverted 35 mm glass bottom microwell petri dish glued onto the sealed plate using Twinsil Speed silicone. The glass bottom of the petri dish is removed so that the chamber has an opening to add water over the sealed wells.

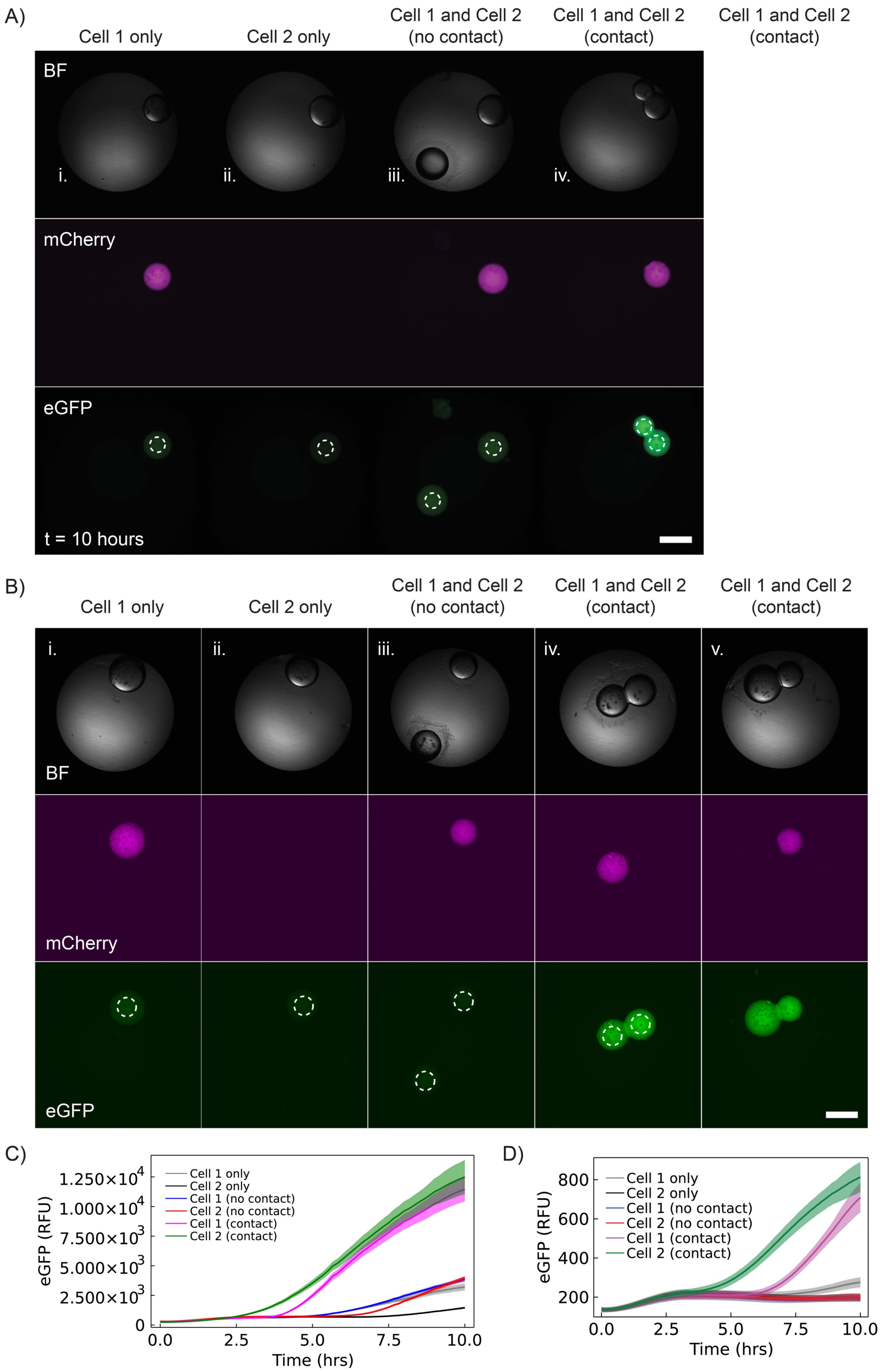


**Figure S11.** (A) Brightfield (BF), mCherry, and eGFP channels of the feedback synthetic cells at endpoint (t = 10 hours). (i) Cell 1 in isolation containing 0.5 nM pT7 LuxI, 0.5 nM pT7 LasR, 10 nM pLas eGFP, and 7.2 µM purified mCherry protein as a marker. (ii) Cell 2 in isolation containing 0.5 nM pT7 LuxR, 10 nM pLux eGFP, and 0.1 nM pLux LasI. (iii) Cell 1 and Cell 2 in the same well but not in contact. (iv) Cell 1 and Cell 2 in contact with each other. Scale bar is 500 µm. (B) Repeat experiment of feedback synthetic cells. Two samples of feedback cells in contact samples are shown in (iv) and (v). (C-D) Timelapse of eGFP RFU signal of the same synthetic cells in A and B from 0-10 hours showing the activation of Cell 2 and the Cell 1 by feedback when in contact. All data was obtained from timelapse widefield fluorescence microscopy using a using a Zeiss Andor Axiovert 200M with a 5x/0.15 Plan-Neofluar Ph1 M27 objective. Samples were incubated at room

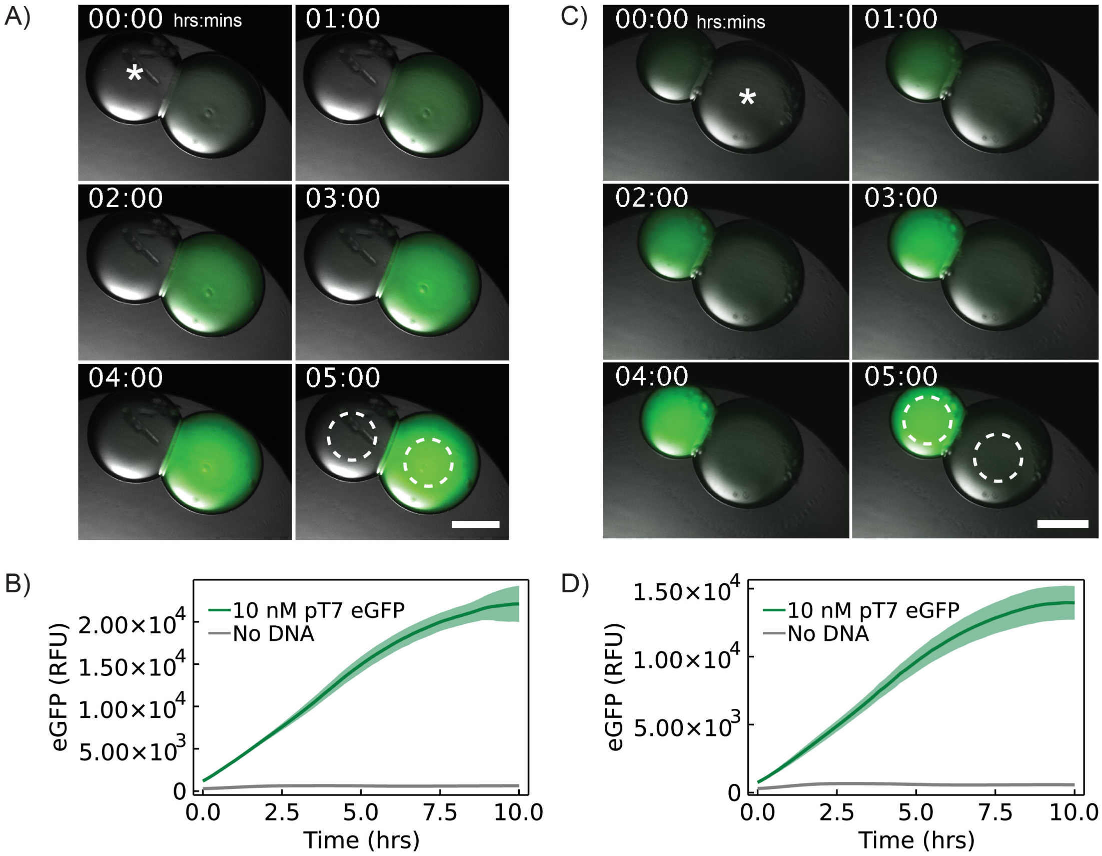


**Figure S12.** eGFP expression in DIB synthetic cells showing that DNA, transcribed mRNA, and translated protein are not permeable through the lipid bilayer interface. (A) Widefield microscopy images of GFP and brightfield images overlaid of a pair of synthetic cells in contact with one another. The cells contain 10 nM pT7 eGFP plasmid and without DNA (marked *) from 0-5 hours. Data shows the same trends as those shown in Fig. 3 in the main text. (B) Plot showing eGFP RFU intensity over time of the two synthetic cells shown in A. Mean (bold line) and standard deviation (shaded region) RFU were taken from the areas denoted in A with dashed lines. (C-D) Replicate experiment of a pair of synthetic cells with 10 nM pT7 eGFP plasmid and without DNA (marked *). Scale bars are all 250 µm. All data was obtained from timelapse widefield fluorescence microscopy using a Zeiss Andor Axiovert 200M with a 5x/0.15 Plan-Neofluar Ph1 M27 objective. Samples were incubated at room temperature (24-27°C).

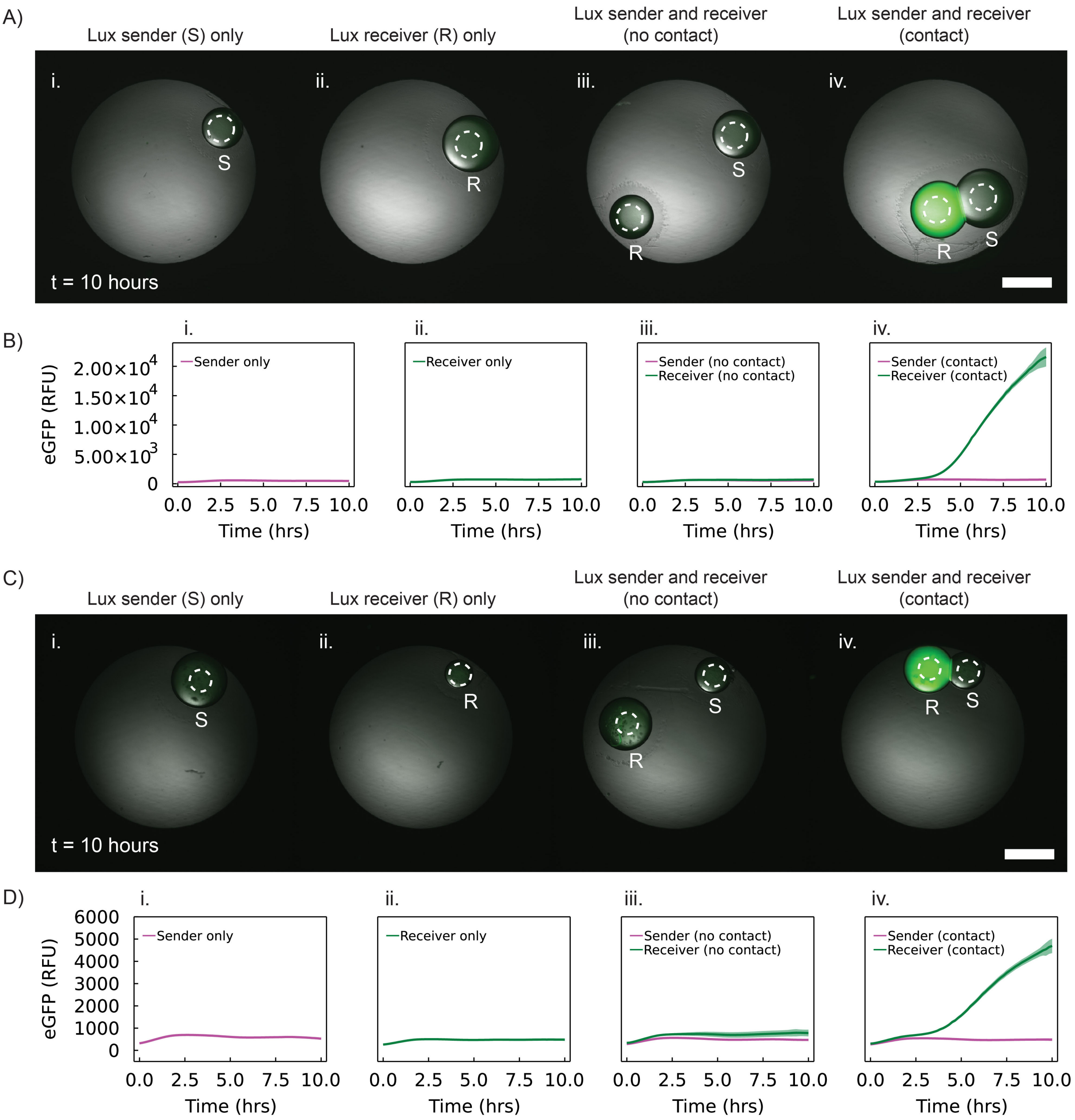


**Figure S13.** (A) Microscopy images showing eGFP fluorescence at t = 6 hours of Lux sender and receiver synthetic cells in isolation (i and ii), within one well but not in contact (iii), and in contact (iv). Images are composed of merged brightfield and GFP fluorescence channels. Scale bar is 500 µm. (B) Plots of eGFP RFU signal as a function of time from the same synthetic cells shown in (A) from 0-10 hours. Mean (solid line) and standard deviation (shaded area) RFU were taken from the areas denoted in A with dashed lines. (C-D) Repeat experiment for Lux sender and receiver cells as A-B. All data was obtained from timelapse widefield fluorescence microscopy using a Zeiss Andor Axiovert 200M with a 5x/0.15 Plan-Neofluar Ph1 M27 objective. Samples were incubated at room temperature (24-27°C).

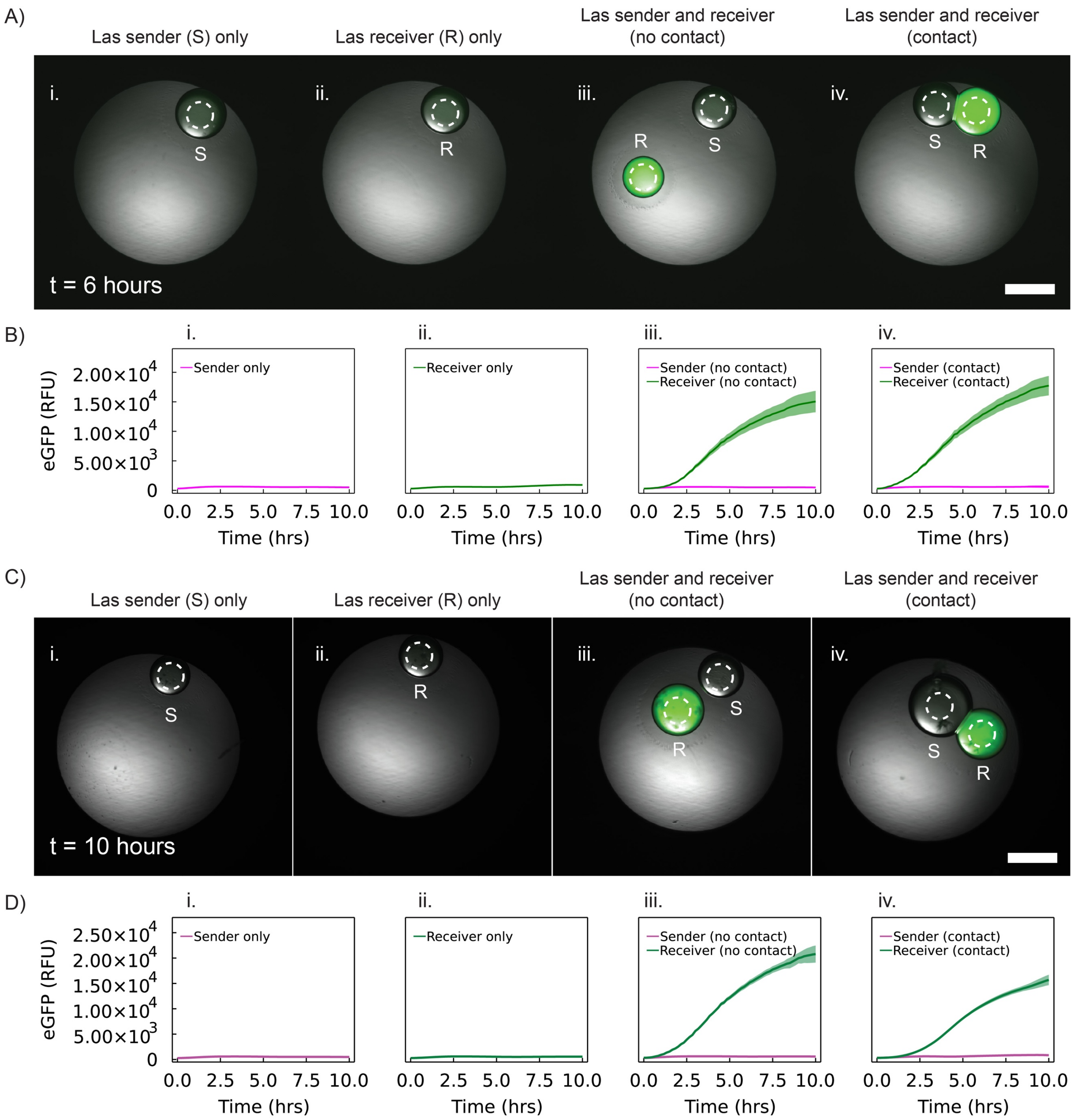


**Figure S14.** (A) Microscopy images at t = 6 hours of Las sender and receiver synthetic cells in isolation (i and ii), within once well but not in contact (iii), and in contact (iv). Images are composed of merged brightfield and GFP fluorescence channels. Scale bar is 500 μm. (B) Plots of eGFP RFU signal as a function of time from the same synthetic cells shown in (A) from 0-10 hours. (C-D) Repeat experiment for Las sender and receiver cells as A-B. All data was obtained from timelapse widefield fluorescence microscopy using a Zeiss Andor Axiovert 200M with a 5x/0.15 Plan-Neofluar Ph1 M27 objective. Samples were incubated at room temperature (24-27°C).

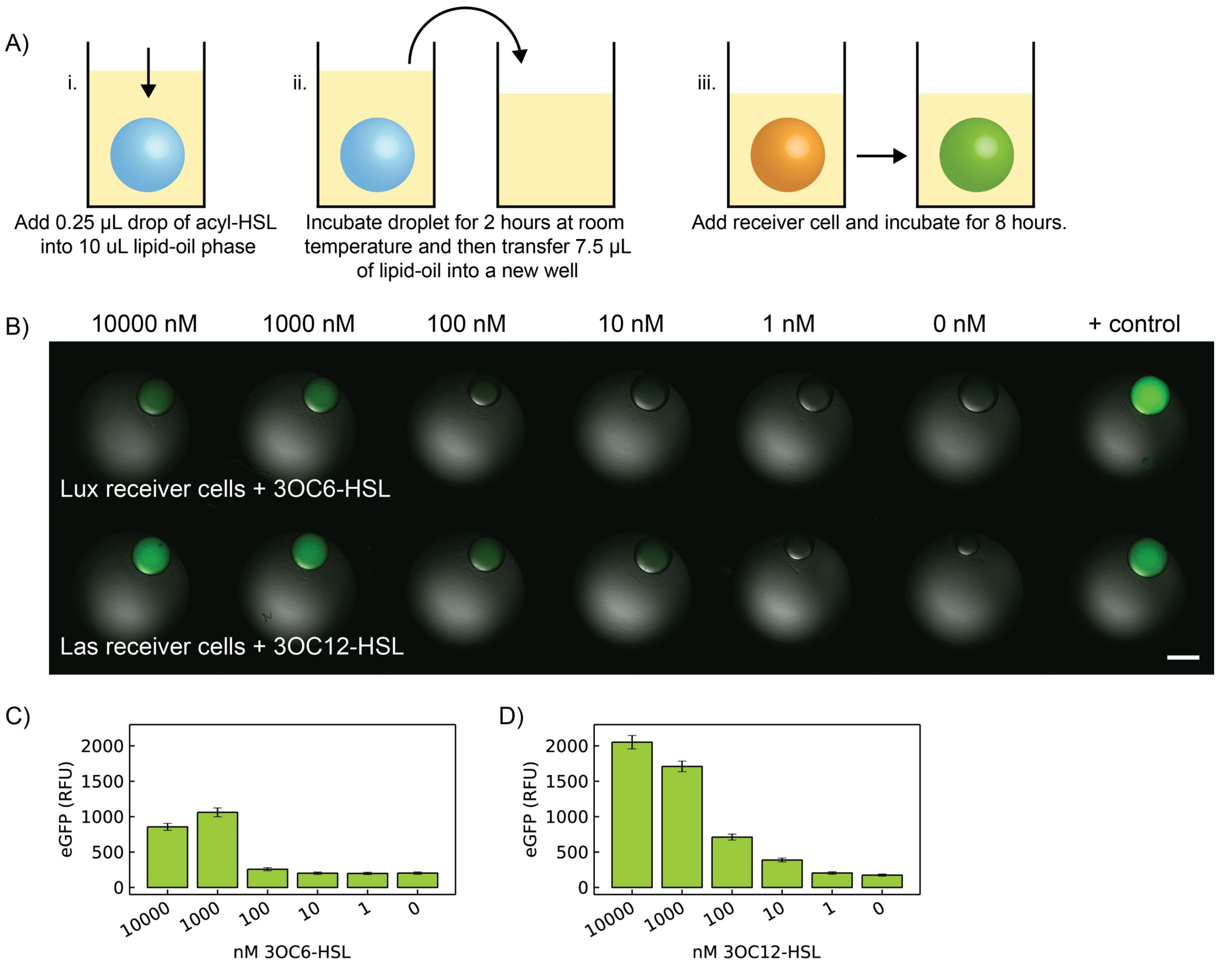


**Figure S15.** (A) Illustration of partitioning assay of acyl-HSL into the oil phase. First, (i) a 0.25 µL drop of the acyl-HSL is added into 10 µL of the lipid-oil phase. (ii) After 2 hours of incubation, 7.5 µL of the oil phase is transferred to a new well without the droplet. (iii) A receiver cell is added into the new 7.5 µL oil phase well, incubated for 8 hours, and imaged for eGFP expression. (B) Endpoint eGFP expression (t = 8 hours) of receiver cells (0.5 nM pT7 LuxR and 10 nM pLux eGFP or 0.5 nM pT7 LasR and 10 nM pLas eGFP) in a lipid-oil phase preincubated with different concentrations of 0.25 µL 3OC6-HSL or 3OC12-HSL droplets. Positive control are the same receiver cells with 1000 nM of the HSL directly added into the CFES mix. Scale bar is 500 µm. (C-D) RFU values quantified from the image in (B) showing that induction of the Lux system begins at the 100 nM 3OC6-HSL preincubated droplet and 10 nM 3OC12-HSL preincubated droplet for the Las system. All data was obtained from timelapse widefield fluorescence microscopy using a using a Zeiss Andor Axiovert 200M with a 5x/0.15 Plan-Neofluar Ph1 M27 objective. Samples were incubated at room temperature (24-27°C).

**S6. Echo-made synthetic cell populations**

Custom synthetic cell populations were made using an Echo 550 acoustic liquid handler to generate droplets between 2.5-20 nL from a source plate containing the CFES mix (prepared as described previously) into the destination plate with 2 uL of lipid-oil phase. To form DIB populations, the destination plate is flipped right-side up and incubated at room temperature for 1 minute between addition of new droplets in the same well to allow DIBs to stabilize their lipid monolayers before coming into contact with other droplets. This is done by assigning a 'new' plate between droplet transfers in the transfer plan to eject the destination plate (Table S12). Populations with different synthetic cell types are made by preparing multiple CFES mixes with different plasmid DNA and loaded into separate wells in the source plate for dispensing (as demonstrated also in Table S12). Synthetic cell populations were prepared with 2.5 nL sized droplets. Constitutive expression of eGFP in these 2.5 nL synthetic cells worked, but the Lux receiver cells were not induced by Lux sender cells (Fig. S16). Preparation of larger DIBS (20 nL) showed positive expression of eGFP within the receiver cells. (Fig. S17).

Heterogeneous populations of gene expressing droplets were prepared. Lux receiver cells (10 nM pLux eGFP and 0.5 nM pT7 LuxR) were combined with Lux quorum sensing (QS) cells (10 nM pLux eGFP, 0.5 nM pT7 LuxI, and 0.5 nM pT7 LuxR) into a heterogeneous synthetic cell population. Individually, the receiver cells did not express eGFP due to the absence of inducer molecule 3OC6-HSL, while the QS cells express eGFP due to the production of 3OC6-HSL synthase from pT7 LuxI. However, when the two cell types are combined into one population, all cells showed positive expression of eGFP due to the diffusing 3OC6-HSL from the QS cells (Fig. S18).

**Table S12**. Echo 550 transfer plan for making four synthetic cells (two synthetic cell types from source well C3 and C4) in well F5. The Destination Barcode for the destination plate is assigned different names to tell the instrument to remove the plate to allow the DIBs to stabilize. However, the same destination plate is returned to add multiple droplets into the same well.

| **Source Barcode** | **Source Well** | **Destination Barcode** | **Destination Well** | **Volume** |
|---|---|---|---|---|
| S1 | C3 | D1 | F5 | 10 |
| S1 | C3 | D2 | F5 | 10 |
| S1 | C4 | D3 | F5 | 10 |
| S1 | C4 | D4 | F5 | 10 |

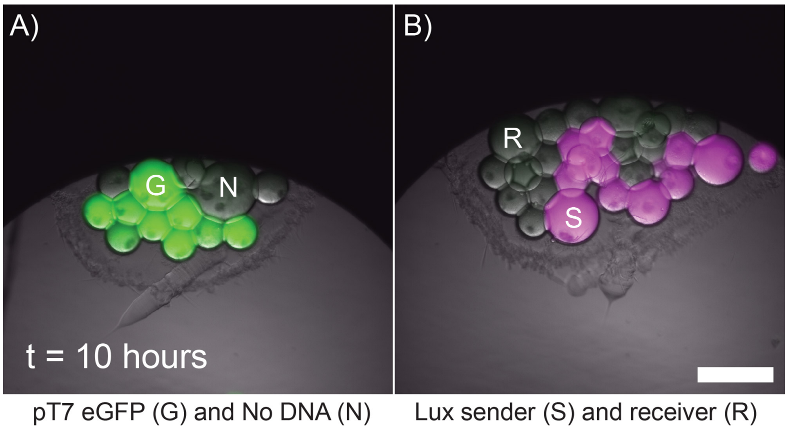


**Figure S16.** (A) Echo-made 2.5 nL synthetic cells with either 10nM pT7 eGFP plasmid (labelled G) or no plasmid DNA (labelled N) after 10 hours of incubation at room temperature. (B) Echo-made 2.5 nL Lux sender cells (0.5 nM pT7 LuxI and 7.2 µM purified mCherry protein as a marker, magenta, labelled S) and receiver cells (10 nM pLux eGFP and 0.5 nM pT7 LuxR, labelled R) after 10 hours of incubation at room temperature. Images are merged from brightfield, eGFP, and mCherry channels. Scale bar is at 250 µm. Data was obtained from widefield fluorescence microscopy using a Zeiss Andor Axiovert 200M with a 10x/0.3 Plan Neo-fluar Ph1 M27 objective. Samples were incubated at room temperature (24-27°C).

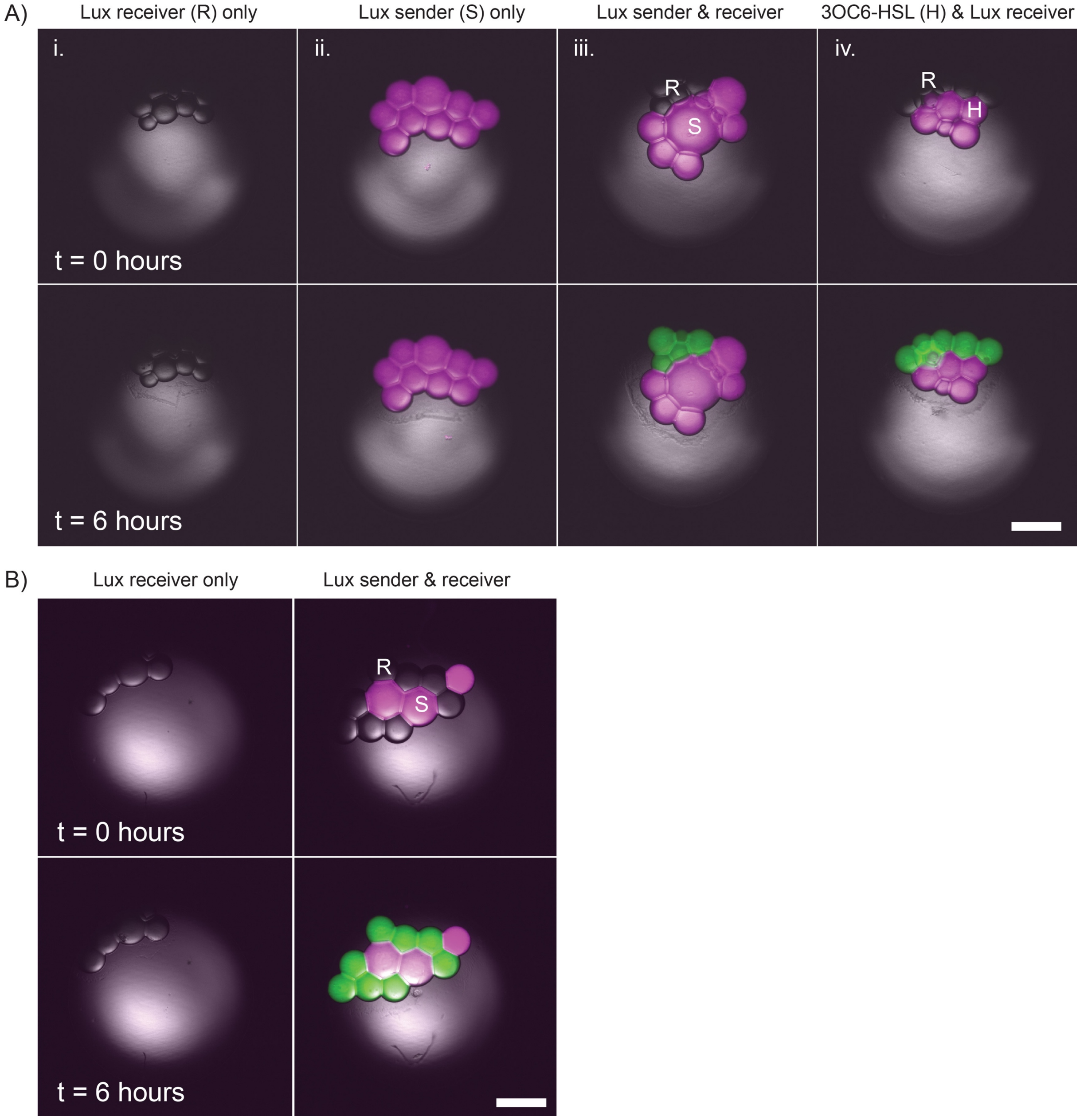


**Figure S17.** Lux sender and receiver synthetic cell populations made by the Echo 550 acoustic liquid handler. (A) 20 nL *E. coli* extract-based CFES droplets of (i) receiver cells only (0.5 nM pT7 LuxR and 10 nM pLux eGFP), (ii) sender cells only (0.5 nM pT7 LuxI) with 7.2 purified µM mCherry protein as marker for the sender cell type, (iii) sender (magenta) and receiver cells (green), and (iv) cells with 1000 nM 3OC6-HSL (magenta) and receiver cells (green) at t = 0 (1st row) and t = 6 hours (2nd row). (B) Replicate experiment of (i) receiver cells only and (ii) sender and receiver cells at t = 0 hours (first row) and t = 6 hours (2nd row). Widefield fluorescence microscopy images are merged from brightfield, eGFP, and mCherry channels. Scale bars are all at 500 µm. All data was obtained from timelapse widefield fluorescence microscopy using a using a Zeiss Andor Axiovert 200M with a 5x/0.15 Plan-Neofluar Ph1 M27 objective. Samples were incubated at room temperature (24-27°C).

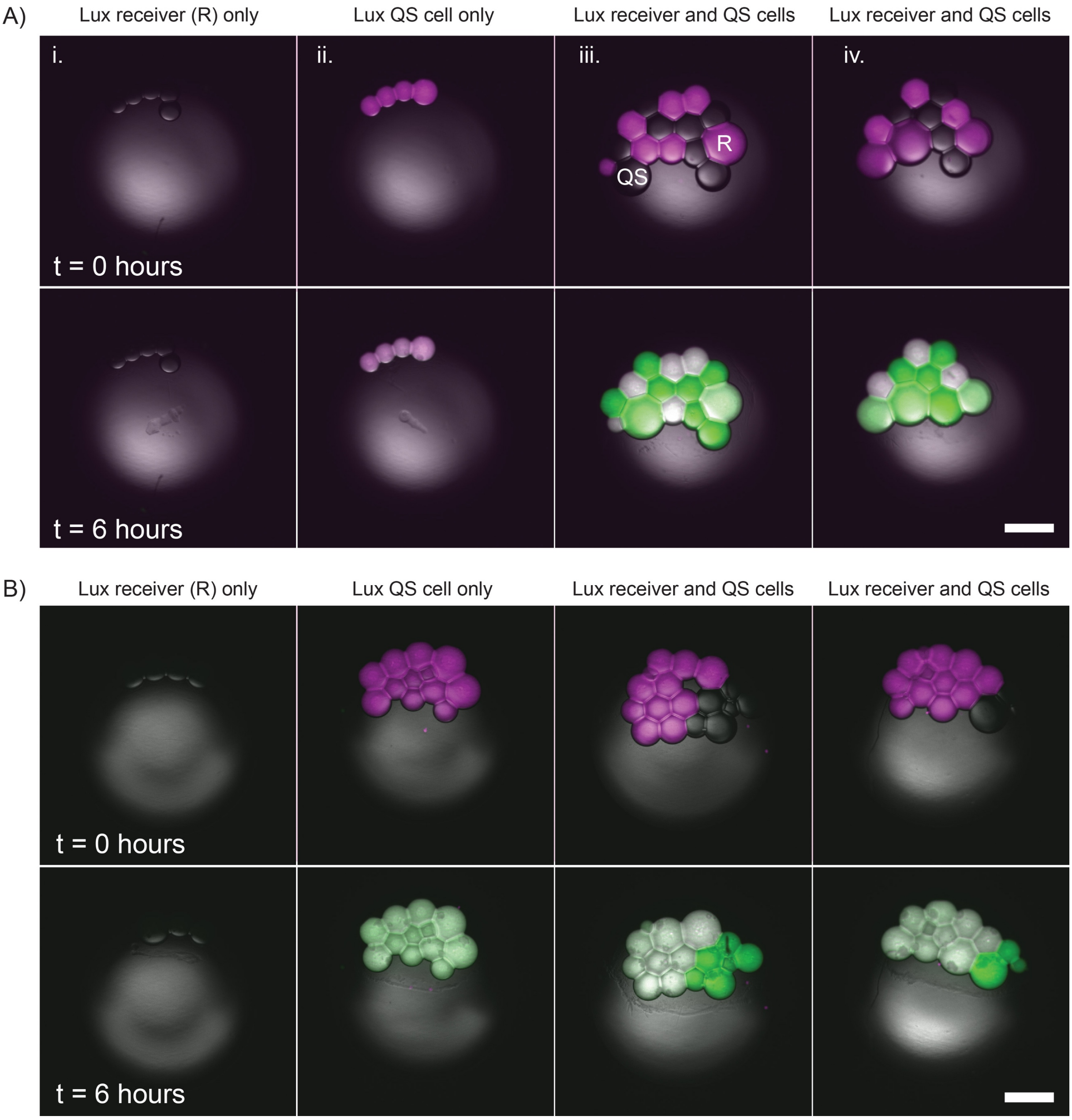


**Figure S18.** Heterogeneous synthetic cell population uniformly expresses eGFP because of quorum sensing communication. (A) 20 nL *E. coli* extract-based CFES droplets of (i) receiver cells only (0.5 nM pT7 LuxR and 10 nM pLux eGFP), (ii) quorum sensing (QS) cells only (0.5 nM pT7 LuxR, 10 nM pLux eGFP, and 0.5 nM pT7 LuxI) with 7.2 purified µM mCherry protein as marker for the QS cell type, (iii-iv) QS (magenta) and receiver cells (green) at t = 0 (1st row) and t = 6 hours (2nd row). (B) Replicate experiment of (i). Widefield fluorescence microscopy images are merged from brightfield, eGFP, and mCherry channels. Scale bars are all at 500 µm. All data was obtained from timelapse widefield fluorescence microscopy using a Zeiss Andor Axiovert 200M with a 5x/0.15 Plan-Neofluar Ph1 M27 objective. Samples were incubated at room temperature (24-27°C).
